# A synergistic transcriptional regulation of olfactory genes derives complex behavioral responses in the mosquito *Anopheles culicifacies*

**DOI:** 10.1101/218081

**Authors:** Tanwee Das De, Tina Thomas, Sonia Verma, Deepak Singla, Charu Rawal, Vartika Srivastava, Punita Sharma, Seena Kumari, Sanjay Tevatiya, Jyoti Rani, Yasha Hasija, Kailash C Pandey, Rajnikant Dixit

## Abstract

Decoding the molecular basis of host seeking and blood feeding behavioral evolution/adaptation in the adult female mosquito may provide an opportunity to design new molecular strategy to disrupt human-mosquito interactions. However, despite the great progress in the field of mosquito olfaction and chemo-detection, little is known that how the sex-specific specialization of the olfactory system enables adult female mosquitoes to derive and manage complex blood feeding associated behavioral responses. A comprehensive RNAseq analysis of prior and post blood meal olfactory system of *An. culicifacies* mosquito revealed that a minor but unique change in the nature and regulation of key olfactory genes play a pivotal role in managing diverse behavioral responses. Age dependent transcriptional profiling demonstrated that adult female mosquito’s chemosensory system gradually learned and matured to drive the host-seeking and blood feeding behavior at the age of 5-6 days. A zeitgeber time scale expression analysis of Odorant Binding Proteins (OBPs) unravels unique association with a late evening to midnight peak biting time. Blood meal-induced switching of unique sets of OBP genes and Odorant Receptors (ORs) expression coincides with the change in the innate physiological status of the mosquitoes. Blood meal follows up experiments provide enough evidence that how a synergistic and concurrent action of OBPs-ORs may drive ‘prior and post blood meal’ complex behavioral events. Finally, tissue-specific gene expression analysis and molecular modelling predicted two uncharacterized novel sensory appendages proteins (SAP-1 & SAP2) unique to *An. culicifacies* mosquito and may play a central role in the host-seeking behavior.

**Significance:** Evolution and adaptation of blood feeding behavior not only favored the reproductive success of adult female mosquito but also make them an important disease vectors. Immediately after emergence, an environmental exposure may favor the broadly tuned olfactory system of mosquitoes to derive complex behavioral responses. But, how these olfactory derived genetic factors manage female specific ‘pre and post’ blood meal associated complex behavioral responses are not well known. We unraveled synergistic actions of olfactory factors governs an innate to prime learning strategy to facilitate rapid blood meal acquisition and downstream behavioral activities. A species-specific transcriptional profiling and an *in-silico* analysis predict novel ‘sensory appendages protein’, as a unique target to design disorientation strategy against the mosquito *Anopheles culicifacies*.

## Introduction

Mosquitoes are one of the deadliest animals in the world, which are responsible for transmitting a variety of infectious disease such as malaria, dengue fever, chikungunya, zika fever worldwide. According to WHO last annual report, malaria is one of the major vector-borne diseases that causes 212 million morbidity cases and more than 4 million mortalities (1). In India, malaria situation is more complex, where WHO states that the socio-economic burden of $1.94 billion due to malaria alone (1). Current tools to control and manage malaria face challenges due to the emergence of parasite resistance to antimalarial drugs and insecticide resistance of mosquitoes (2), (3), (4), (5), (6), (7). Thus, alternative molecular tools are needed to rule out the expanding vector population as well as parasite development.

One of the key molecular strategies under not-to-bite approach relies on the designing of a new class of molecular tools able to disorient/alter the adult female mosquito host-seeking behavior (8). However, defining the molecular basis of host-seeking behavioral evolution and adaption to blood feeding by the adult female mosquitoes remains central to our understanding. Probably this may be due to the complex interaction of genetic and non-genetic environmental factors deriving mosquito navigation (9). In nature mosquitoes encounter many challenges to sustain in daily life i.e. they rely immensely on their sense of smell (olfaction) for the majority of their lifecycle stages (8). Mosquitoes are able to detect and discriminate thousands of odor molecules, which may further vary depending on the host preference. These complex behavioral events are largely mediated by the diverse chemosensory genes encoding odorant binding proteins (OBPs), odorant degrading enzymes (ODEs), odorant receptors (ORs) and other accessory proteins including sensory neuron membrane protein (SNMP) (10). Odorant binding proteins (OBPs), which are bathed within the sensillum lymph, are low molecular weight soluble proteins that mediate the first interaction of the olfactory system with the external world (10), (11), (12). These globular protein molecules showed significant diversity within the same family and are believed to bind with a wide range of hydrophobic odorant molecule. After binding with these odor molecules, OBPs shuttle it to their respective olfactory receptors present on olfactory receptor neurons (ORNs) (13), (10), (14). Insect olfactory receptors (OrX) are present to work in association with the obligate receptor co-receptor (Orco) on the dendritic membrane of ORN (10). Orco is essential for proper dendritic trafficking of the OrX, facilitating the formation of odorant gated ion channels by structural alteration that is opened upon odorant binding (10), (15). Thus, it is plausible to hypothesize that prior blood meal, key interactions of odorants and their cognate receptors may have a significant influence on food choice decision and blood meal uptake process.

For a successful blood feeding event, an adult female mosquito need to negotiate and manage multiple behavioral coordinates including searching, locating, landing over a suitable host, followed by tracing the proper site to pierce and suck the blood within two minutes (15), (16), (17), (18), (19), (20) (21). Just after the piercing organ (proboscis), it is the salivary glands which mediate the immediate biochemical interaction with the vertebrate blood and facilitate rapid blood meal uptake. Our recent study suggested that adult female mosquito’s salivary glands are evolved with the unique ability of gene expression switching to manage meal specific (sugar vs blood) responses (22), but the molecular nature of the olfactory and neuro-system in regulating the salivary gland function is yet to unravel.

Post blood meal mosquitoes need to enter into a new habitat favouring successful oviposition (23), (24). In fact, after blood meal acquisition, mosquitoes undergo two major behavioral switching events (i) search suitable site(s) for temporary resting and completion of blood meal digestion (~30hrs) which is necessary for egg maturation (48-72hrs); and (ii) find proper oviposition site for successful egg laying (25). After completion of egg laying event, the adult female mosquitoes regain host-seeking behavioral activities for a second blood meal to complete the next gonotrophic cycles (24), (26). Notably, ‘prior and post’ blood meal associated habitats may have a significant difference in their physical, chemical and biological characteristics (23), but how olfactory driven factors manages these complex events is still an unresolved puzzle (27).

Immediately after emergence an exposure to diverse environmental/chemical cues facilitate olfactory maturation and learning for different innate behavioral activities such as feeding and mating in both the sexes (9). However, in case of adult female mosquitoes, we opined that the evolutionary forces, might have favoured an extra but unique specialization of host seeking and blood feeding behavioral adaptation, possibly a result of frequent mosquito-vertebrate interactions **(Fig.1).** Once, a mosquito takes first blood meal it needs to manage major physiological activities linked to blood meal digestion and egg maturation. These physiological changes possibly may have another level of impact on olfactory responses to guide oviposition site finding behavior. We further hypothesize that first blood meal exposure must have a priming effect on the olfactory responses expediting the consecutive host seeking and blood feeding behavioral activities more rapidly than previous one.

**Figure 1:**
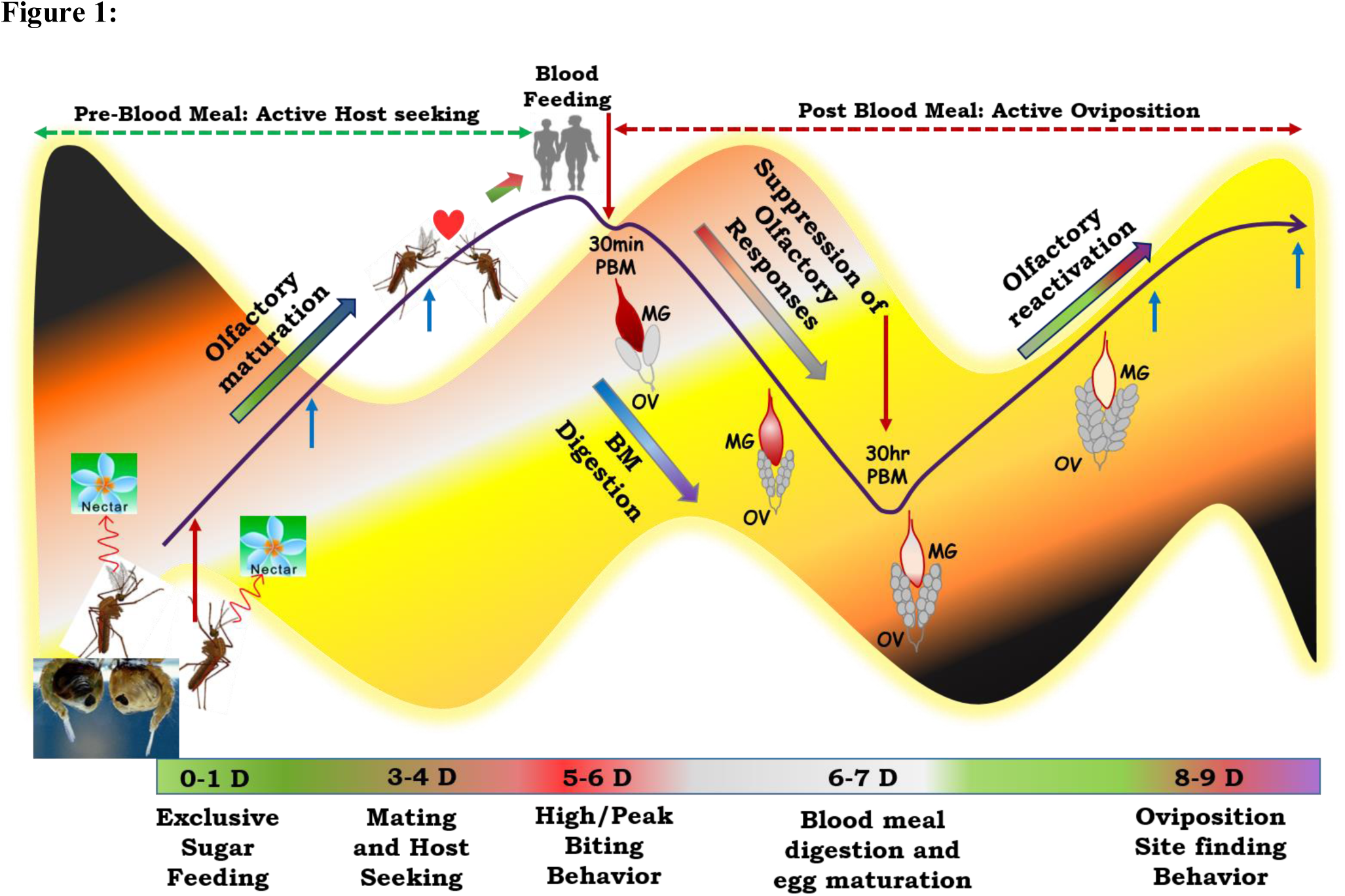
Working Hypothesis to establish functional co-relation of the olfactory system under distinct feeding status. Mosquitoes Adult mosquitoes, just after emergence from pupae are exclusive sugar feeders and dependent on nectar sugar to acquire energy for flight activity. Exposure of the adult mosquitoes to the diverse aromatic environment facilitate their learning and maturation of the olfactory system which enable successful mating and host seeking behavioral activities. But the function of the olfactory system starts to diminish just after blood feeding and become ceased at least for 30hrs of post blood meal. Blood feeding initiates lots of physiological changes including blood meal digestion in the midgut and egg maturation in the ovary which consume lots of energy and thus mosquitoes manipulate the energy cost by shutting down the olfactory responses and preferred to take rest at a cool dark place. After 30 hrs of blood feeding the blood almost digested in the midgut and maturation of egg reached a threshold level which reinforce the mosquito to perform to next level of behavior. Thus, recovery/reactivation of the olfactory responses occur to find a suitable site for egg laying/oviposition. To capture these molecular snapshot and track the events, we collected olfactory tissues at three different physiological conditions for RNASeq analysis (Highlighted as red arrows), and coupled with gene expression study with more elaborated time and physiological state (highlighted with blue arrows). MG: Midgut; OV: Ovary. Mosquitoes each and every life cycle stages are tightly regulated by circadian (dawn & dusk) cycle (Background light dark color code).

To test and decode this evolutionary speciality, we performed RNAseq transcriptomic analysis of the olfactory system of adult female *An. culicifacies* mosquito, responsible for more than 65% malaria cases in India (28). A comprehensive molecular and functional annotation of RNAseq data unravelled a limited but remarkable change in the nature and regulation of unique sets of olfactory gene repertoire in response to distinct feeding status of the mosquitoes. Extensive transcriptional profiling of the selected transcripts showed biphasic and synergistic regulation under the distinct innate physiological status of the mosquitoes, possibly to facilitate and manage the complex host-seeking behavioral events. Finally, our structural bioinformatic analysis predicts the key residues of the selected sensory appendages proteins, targets for future functional validation and characterization.

## Material and Methods

Fig. 2 represents a technical overview of the current investigation.

**Figure 2:**
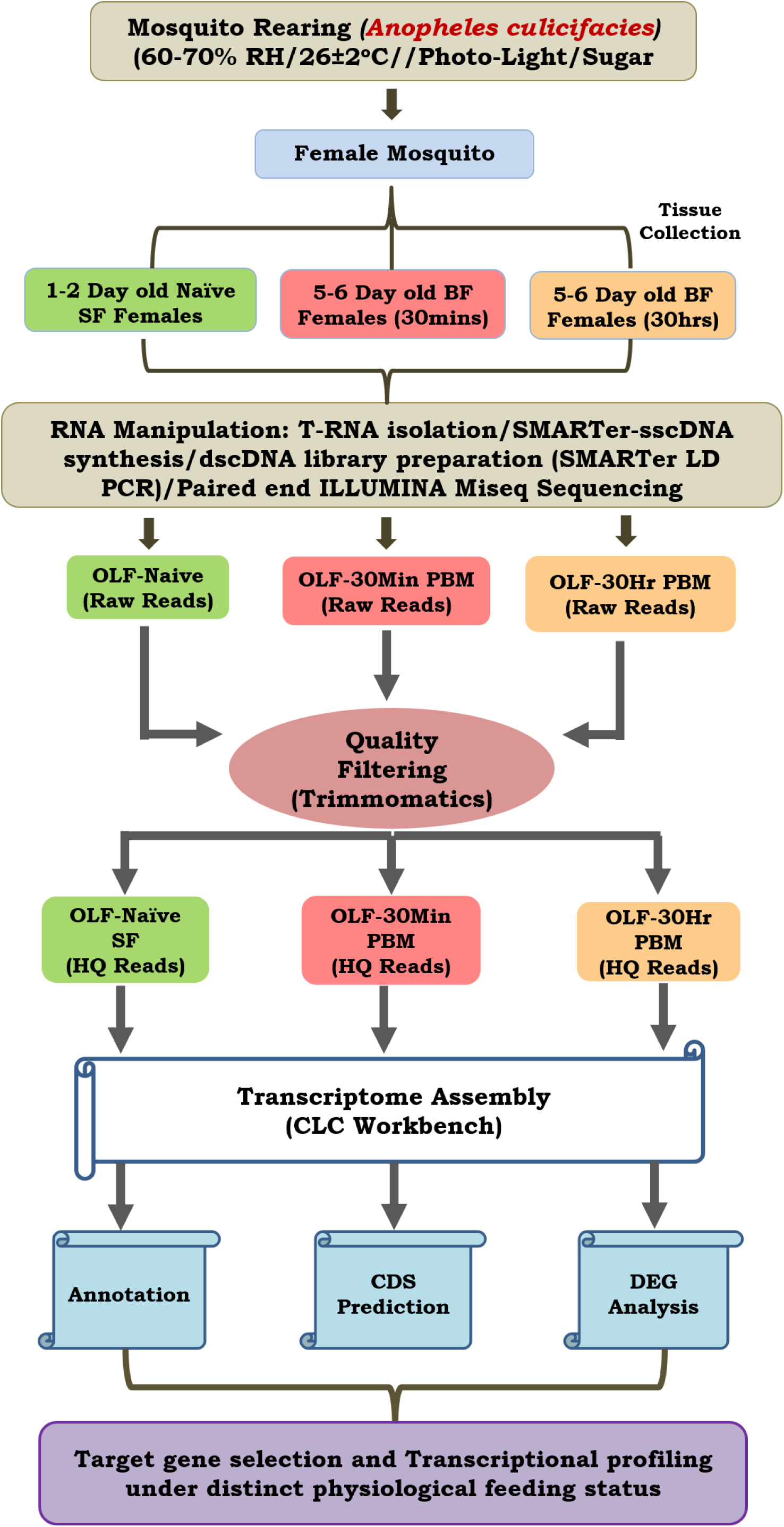
A technical overview to decode the hard-wired genetic structure of olfactory system of *Anopheles culicifacies*. We sequenced and analysed three RNA samples extracted from the olfactory tissues of approximately 30 mosquitoes individually and pooled to form one single sample. With this strategy we were able to quantify and validate the estimation of gene expression, expected with a minimum chance of aberrations as described earlier (21).

**Mosquito Rearing and Maintenance:** A cyclic colony of the mosquito *An. culicifacies*, sibling species A and *An. stephensi* were reared and maintained at 28±2°C, RH=80% in the central insectary facility as mentioned previously (22), (29). All protocols for rearing and maintenance of the mosquito culture were approved by ethical committee of the institute.

**RNA isolation and Transcriptome Sequencing Analysis:** Complete olfactory tissue which includes antennae, maxillary palp, proboscis and labium, were dissected from 0-1 day of age, 30 min post blood fed and 30 hrs post blood fed *An. culicifacies* mosquito and collected in Trizol Reagent. Total RNA was isolated and double-stranded cDNA library was prepared by a well-established PCR-based protocol described previously (22), (30). Whole transcriptome sequencing of the olfactory tissue was performed by following the Illumina MiSeq 2 × 150 paired-end library preparation protocol. The sequencing data analysis pipeline is shown in Fig 2. A *denovo* clustering using CLC Genomics workbench (V6.2) (31) was used to build final contigs/transcripts dataset for functional annotation. For comprehensive annotation and differential gene expression analysis we followed essentially the same protocol as described earlier (22). Relative quantification of the genes was calculated by FPKM values (Fragments Per Kilobase of Exon Per Million Fragments Mapped).

**Identification and molecular cataloguing of olfactory genes in *An. culicifacies*:** An initial BLAST search analysis predicted a total of 93 transcripts encoding putative OBP homologs from the olfactory transcriptome data of *An. culicifacies* mosquito. To predict additional OBPs, a merged database of mosquito and *Drosophila* OBPs was re-queried against *An. culicifacies* draft genome/predicted transcripts databases available at <u>www.vectorbase.org</u> and build up the final OBP catalogue and classification for phylogenetic analysis as detailed in the supplemental **figure S1a**. A PDB database homology search analysis and GO annotation was used to identify and catalogue other putative olfactory receptor genes manually.

**PCR based Gene Expression Analysis:** The head tissue containing the olfactory appendages of female *An. culicifacies* mosquito was dissected at different zeitgeber time point. The 24 hr time scale of the LD cycle is represented as different Zeitgeber time (ZT) where ZT0 indicate the end of dawn transition, ZT11 is defined as the start of the dusk transition and ZT12 is defined as the time of lights off (32). Along with that, different tissues viz. head (male, female), Legs (male, female), brain, olfactory tissue (OLF), female reproductive organ (FRO) and Male reproductive organ (MRO) of both *An. culicifacies* and *An. stephensi* mosquitoes were dissected and collected in Trizol followed by total RNA extraction and cDNA preparation. Differential gene expression analysis was performed using the normal RT-PCR and agarose gel electrophoresis protocol. For relative gene expression analysis, SYBR green qPCR (Thermo Scientific) master mix and Illumina Eco Real-Time PCR machine were used. PCR cycle parameters involved an initial denaturation at 95^o^C for 5 min, 40 cycles of 10 s at 95^o^C, 15 s at 52^o^C, and 22 s at 72^o^C. Fluorescence readings were taken at 72^o^C after each cycle. The final steps of PCR at 95^o^C for 15 sec followed by 55^o^C for 15 sec and again 95^o^C for 15 sec was completed before deriving a melting curve. To better evaluate the relative expression, each experiment was performed in three independent biological replicates. Actin or S7 gene were used as internal control in all the experiment and the relative quantification was analysed by 2^−ΔΔCt^ method (33). Differential gene expression was statistically analysed using student ‘t’ test.

**Blood meal time series follow up:** For blood meal time series follow up the experiment, the olfactory tissues were collected from both naïve sugar fed and blood fed mosquitoes at different time points. Olfactory tissues collections were initiated from 0-1 day of naïve sugar-fed mosquitoes and proceed up to 6-7 days on every alternative day. After the 6^th^ day, the adult female mosquitoes were offered first blood meal by offering a live animal (rabbit) and immediately collected olfactory tissues for 30 minutes’ time point. The full blood-fed mosquitoes were separated and kept in a proper insectary condition for further experiment. After collection of olfactory Tissues at 30hrs and 72 hrs post blood fed the gravid females were kept for oviposition and again dissected OLF tissues after 24 hrs of the egg laying event. Second blood meal was provided to the egg laid mosquitoes and final collection of OLF tissues were done after 30hrs of 2^nd^ blood meal.

**Structural Modelling of SAP1 and SAP2:** For structure prediction analysis of SAP1 and SAP2 proteins from *An. culicifacies,* initially, template for each query proteins was searched against PDB database using blastp algorithm. Two best templates were selected for each used query sequence and thereafter, modeller9 v.13 was used for the building of 50 models for each query sequence using multiple templates. The best model was selected based on the PROCHECK analysis, and DOPE score. Finally, the selected models were used for binding site prediction using COACH software.

## Results and Discussion

It is well known that a circadian dependent modulation of olfactory responses significantly influences the biting behavior in *Anopheline* mosquito species (32). But the knowledge that how this olfactory derived modulation manages pre and post blood meal behavioral events is yet not well understood (24), (25) (27). Based on available literature and knowledge gaps, we argued that immediately after emergence adult female mosquitoes must undergoes two unique changes in their olfactory responses i.e. (i) an exposure to the diverse chemical environment affecting host-seeking behavioral maturation that facilitate feeding preference switching from nectar to blood meal; (ii) blood meal digestion completion, enabling successful egg maturation and re-switching olfactory actions towards oviposition site finding behavior.

To decode and establish this molecular relationship managing ‘prior and post’ blood meal behavioral events we developed a working hypothesis **(Fig. 1**), a plausible mechanism which may have a significant influence on mosquito feeding and survival in diverse ecologies. To test the above hypothesis, first, we generated and analysed a total of ~122 million RNAseq reads of the olfactory tissues collected from 1-2 day old naive (Nv), 5-6-day old immediate blood fed (30m-2h PBM) and 30hr post blood fed (30hr PBM) mosquitoes (Table-1). We chose 30h PBM as a critical time when completion of blood meal digestion occurs in the midgut, which may have direct influence on the reactivation of the olfactory system **(Fig. S2)** (24), (25), (34). For molecular and functional annotation, we assembled each transcriptomic database into contigs/Transcripts and compared against multiple molecular databases as described earlier (22). **Table-S1** represents details of the annotation kinetics of mosquito olfactory databases.

**Blood meal causes modest but unique changes to olfactory responses:** To test whether blood meal alters the global expression pattern of the olfactory transcriptome, we performed a digital gene expression analysis. Initial attempt of cleaned reads mapping to the available draft reference genome failed to yield quality results, probably due to poor annotation (**Fig. S3**). Alternatively, we mapped all the high quality reads against *Denovo* assembled reference map, as described earlier (22). Blood meal causes a modest shift in the transcriptome expression **(Fig. 3a)**, supporting the previous report that first blood meal enhances odorant receptor transcripts abundance modestly, but causes general reduction of mosquito antennal chemosensory gene repertoire in *An. gambiae* (24).

**Figure 3:**
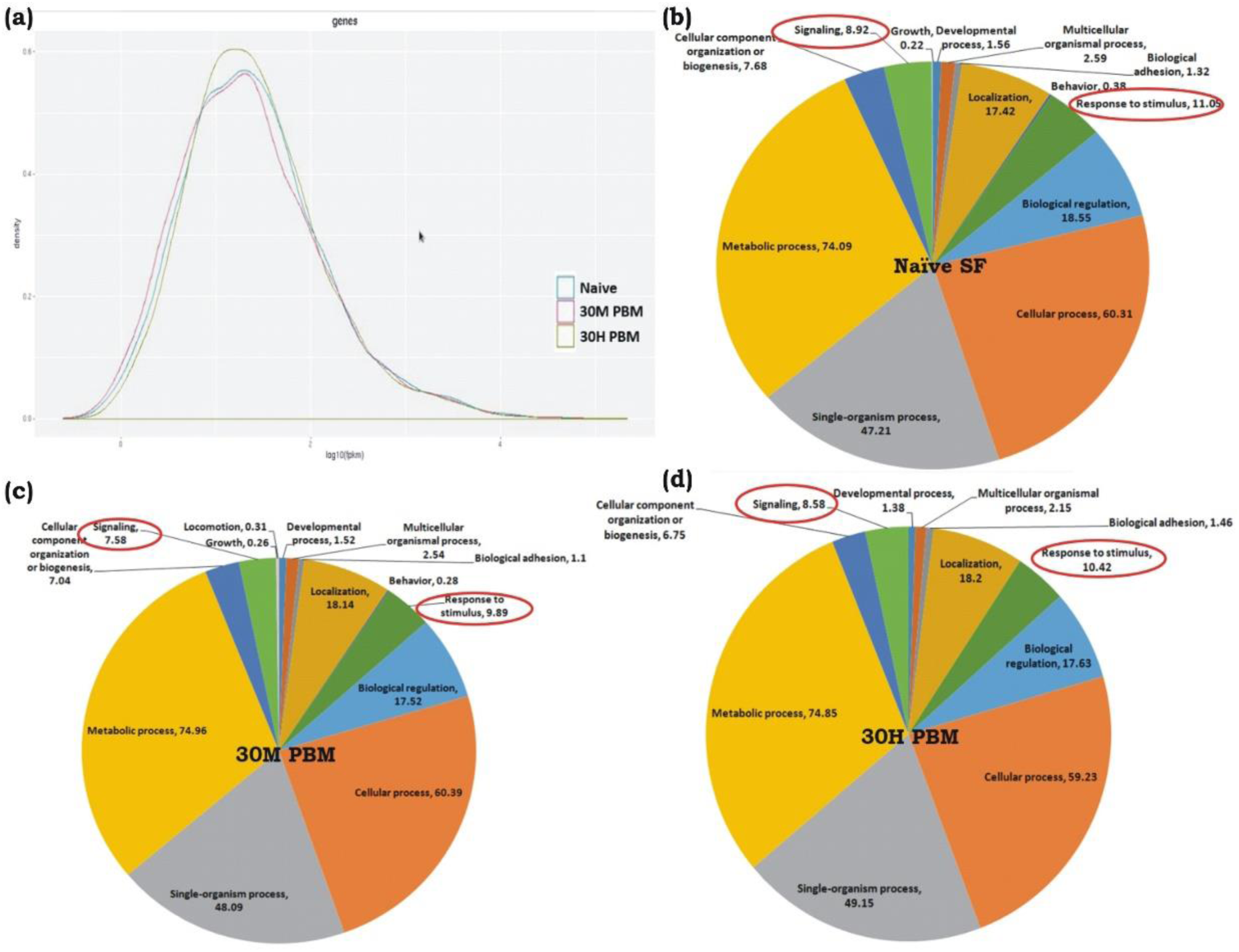
Blood meal cause modest changes in the molecular architecture of the mosquito olfactory system: **(A)** Read density map of the compared Naive; 30M and 30Hr post blood meal (PBM) transcriptomic data of olfactory system; **(B-D)** Functional annotation and molecular catalogue of olfactory transcriptome (Biological Process/Level2/% Transcripts). Red circle marks the genes selected for transcriptional response monitoring. (See Text).

We observed that at least >85% transcriptome remains unaltered, while only ~6% transcripts are up-regulated and ~8.7% transcripts downregulated in 30 min post blood fed samples (**Table-S2 and Dataset S1**). As expected ~10% transcripts expression was further reduced in 30h post blood fed olfactory tissue samples while only 2% transcripts up-regulated when compared to naive unfed mosquitoes (**Table S2**). Interestingly, a comprehensive annotation analysis also predicted that basic composition of the mosquito olfactory tissue does not alter significantly (**Fig. 3b-d**). Owing to a limited change in the olfactory responses, we hypothesize that blood-feeding may not directly cause a major shift in transcript abundance but may alter the functional nature/regulation of the unique transcripts controlling key biological processes such as response to stimulus, circadian rhythm, and signalling in the blood fed adult female mosquitoes. To clarify this complexity, we manually shortlisted the olfactory transcripts either based on their FPKM abundance and/or predicted coding nature and analysed a set of unique genes likely to influence mosquito host-seeking and blood-feeding behavior. To trace the possible molecular link, we extensively profiled their transcriptional regulation under distinct feeding status (see below).

**Daily rhythm and expression change of Odorant binding proteins (OBPs) may influence olfactory responses:** To negotiate and manage the navigation trajectory towards the vertebrate host, olfactory encoded odorant binding proteins (OBPs) play a crucial role to bind and deliver the odorants/chemicals to their cognate odorant receptors, an event guiding behavioral decisions. To explore the possible role of OBPs in the regulation of the olfactory behavior we identified and catalogued a total of sixty-three OBP genes from the mosquito *An. culicifacies* (**Table 1a**). Domain prediction analysis classified the OBPs as Classic OBPs, Plus-C OBPs, Two-domain OBPs and other Chemosensory protein family (**Table 1b; details in table S3**), as described earlier for the mosquito *An. gambiae* (35).

**Table 1.**

| SI No. | Sample Name | Number of OBPs Transcript Retrieved |
| --- | --- | --- |
| 1. | Ac-OLF-Naïve | 14 |
| 2. | Ac-OLF-30min PBM | 10 |
| 3. | Ac-OLF-30Hr PBM | 12 |
| 4. | Ac-Genome Predicted | 27 |
|  | Total | 63 |

(b)
| SI No. | Family of OBPs | Number |
| --- | --- | --- |
| 1. | Classic OBPs | 26 |
| 2. | Plus-C OBPs | 13 |
| 3. | Two-Domain OBPs | 13 |
| 4. | Other Chemosensory Proteins | 11 |

A comprehensive phylogenomic analysis of the Classic putative OBPs of *An. culicifacies* highlights the conserved sequence relationship with *An. gambiae* and other mosquito/insect species (**Fig S4a**), however, Plus-C and Atypical class of OBPs showed a distant relationship to the non-mosquito species *Drosophila* (**Fig. S4b,c**), probably evolved to facilitate host seeking and blood feeding behavior in adult female mosquitoes.

Interestingly, differential gene expression (DGE) data indicated that blood meal restricted the expression of common OBP transcripts (**Fig. S5**). However, first blood meal causes the appearance of unique OBP transcripts (**Table 1**), a crucial event in modulating the behavioral activities in response to changing the feeding status i.e. naive sugar to blood feeding. To further validate and unravel this unique relationship of OBPs regulation, we examined the expression of at least 15 putative OBP transcripts under distinct feeding status of the mosquitoes. In this analysis, we also included two uncharacterized chemosensory proteins (CSPs) named sensory appendage protein (SAP1 & SAP2) having a dominant expression in the naive mosquito olfactory tissue (**Table S3**).

Our Zeitgeber time scale expression showed that out of selected nine OBPs transcripts, at least 6 OBPs showed a >2fold modulation in their expression during late evening to midnight, in the 6-day old naïve mosquitoes **(Fig. 4a).** These data also corroborate with the previous observation of the natural active biting behavior of *An. culicifacies* mosquito in the mid night (36), (37). Surprisingly, sensory appendage proteins (Ac-SAP1 & Ac-SAP2) showed unequivocally an enriched [16 fold for SAP1, p *≤* 0.001 and 6 fold SAP2, p *≤* 0.0001) expression than other tested OBPs. Apart from that, we also observed a transient suppression (30min) and rapid recovery of OBPs expression just after a first blood meal (**Fig. 4b**). Together these observations suggested that a midnight hyper activities of OBPs are able to derive female specific host seeking behavioral activities of *An. culicifacies*. However, a transient change in expression in response to first blood meal further raises a question that how mosquitoes manage blood feeding associated complex behavioral responses such as egg maturation, oviposition etc. We hypothesize those harmonious actions of OBPs with ORs, which are involved in downstream odorant signal transduction cascade, may influence behavioral switching responses from naive sugar feeding status to blood feeding.

**Figure 4:**
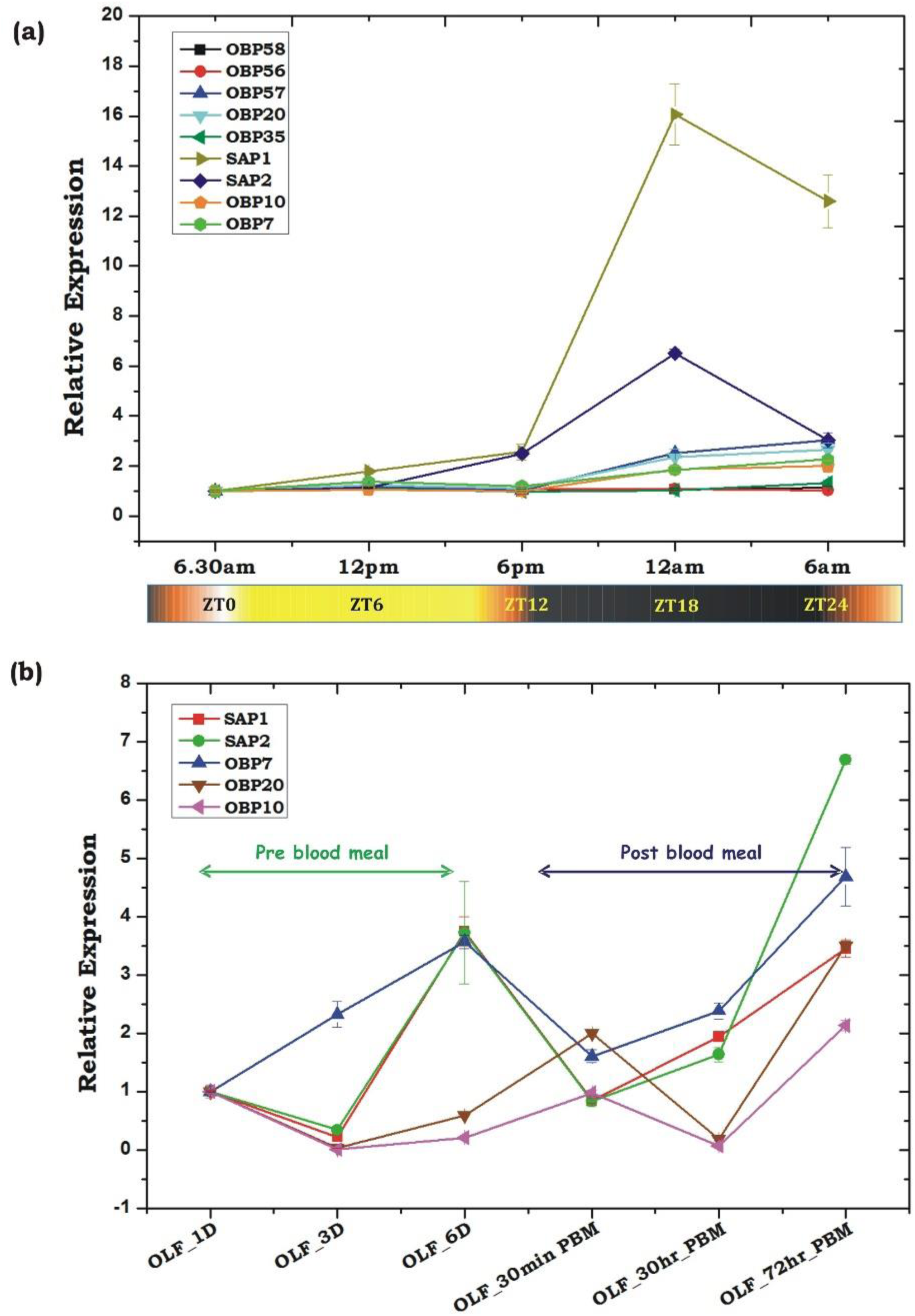
Transcriptional profiling of the odorant binding protein genes (OBPs) under different circumstances. (a) Rhythmic expression of OBP genes in the adult female’s olfactory tissues (OLF) according to different zeitgeber time (ZT) scale, where ZT0 indicate the end of dawn transition, ZT11 is defined as the start of the dusk transition and ZT12 is defined as the time of lights off. (b) Relative expression profiling of OBP genes in pre and post blood fed olfactory tissues. Olfactory tissues (OLF) were collected from 1day, 3day and 6day old sugar fed mosquitoes which were then provided blood meal and then the olfactory tissues were collected after 30mins of post blood fed and 30hrs and 72hrs of post blood fed mosquitoes. The significance of suppression of OBP genes expression after 30hrs of post blood meal are as follows: SAP – ≤ 0.004; SAP2 – ≤ 0.039; OBP7 – ≤ 0.007; OBP20 – ≤ 0.0004; OBP10 – ≤ 0.003.

**Innate physiological status may influence olfactory receptor responses to manage behavioral switching events:** Current literature suggested that a combinatorial coding mechanism of the olfactory receptors enables insects to recognize thousands of diverse chemical cues for selective neuro-actions to meet specific behavioral demands (11), (14), (38). But, the molecular basis that how olfactory receptors superintend and co-ordinate between innate and primed/adaptive odor responses involving ‘prior and post’ blood meal behavioral complexity is yet to be unravelled.

After a successful blood meal, the gut physiology of the naive adult female mosquito undergoes a complex modulation to digest the blood meal and maturation of the eggs. Once the blood meal digestion completed, the mosquitoes may re-switch their olfactory responses for ovposition site finding behavior (24), (39), (40), (41). A transient modulation of OBPs expression in response to blood meal further prompted us to decode and establish its correlation with the olfactory receptors. To unravel this relationship, initially we retrieved, pooled and catalogued a total of 603 unique transcripts linked to response to stimulus and signalling (RTSS) categories **(Fig. 3b,c,d),** encoding diverse nature of proteins such as anion binding, nucleic acid binding, receptor activity, hydrolases and transferase activity (**Fig. 5a**). A comparative GO score distribution analysis predicted lower score hits for the blood-fed cohorts than naive mosquitoes (**Fig. 5a**). Surprisingly, out of 603 transcripts, we noticed only 110 transcripts common to all, while >100 transcripts remain uniquely associated with individual physiological conditions compared in the study (**Fig. 5b**). Together these data suggested that blood meal not only delimits the expression of many olfactory genes but also enriches the expression of many unique transcripts having similar functions.

**Figure 5:**
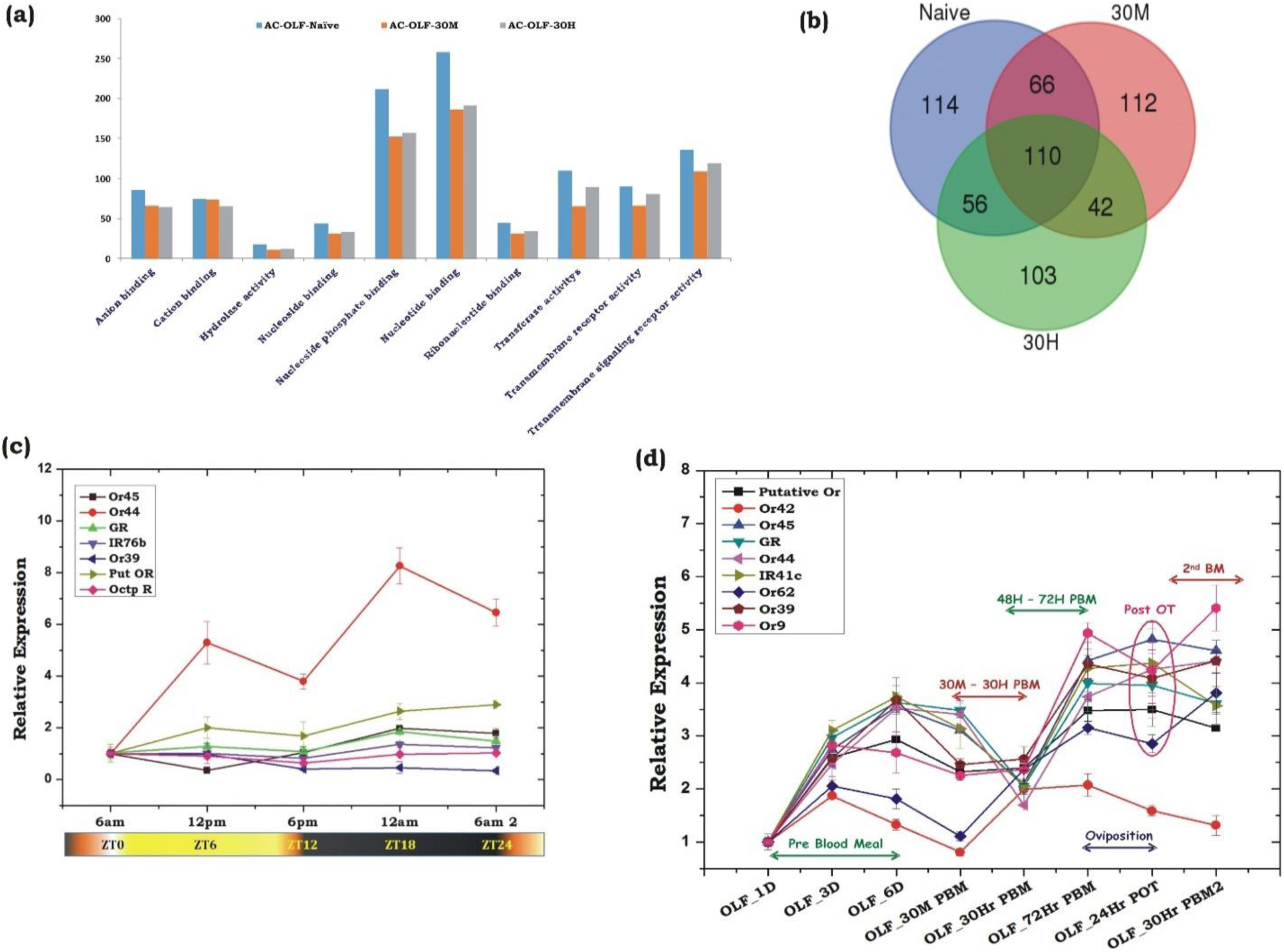
Blood meal modulates odorant receptors expression. **a)** A comparative GO score distribution analysis of response to stimulus and signalling transcripts of naïve and blood fed mosquitoes. b) Venn diagram showing common and unique transcripts of response to stimulus and signalling GO category of naïve and blood fed mosquitoes. (c) Rhythmic expression of olfactory receptor genes (Ors) of *An. culicifacies* in the olfactory tissues of female mosquitoes, where ZT0 indicate the end of dawn transition, ZT11 is defined as the start of the dusk transition and ZT12 is defined as the time of lights off. (d) Transcriptional response of olfactory receptor genes according to blood meal time series experiment. Olfactory tissues (OLF) were collected from naïve sugar fed adult female mosquitoes till 6^th^ day (OLF-1D, OLF-3D, OLF-6D). Then mosquitoes were provided blood meal and again olfactory tissues were collected at different time point after blood feeding, viz. OLF-30M: 30 min post blood fed (PBM); 30hr-PBM: 30hrs of PBM; 72Hr-PBM: 72hrs of post blood meal; then the mosquitoes were kept for oviposition (egg laying), and again the olfactory tissues wer collected 24hrs of post oviposition (24hr-POT). Finally, the 2^nd^ blood meals were provided to the egg laid mosquitoes and collected olfactory tissue 30hrs of second blood meal (30hr-PBM2). The significance of suppression of OR genes expression after 30hrs of post blood meal are as follows: Putative Or - ≤ 0.001; Or42 – ≤ 0.05; GR – ≤ 0.003; Or44 – ≤ 0.0002; IR41c – ≤ 3.5E^−05^; Or62 – ≤ 0.06; Or39 – ≤ 0.02; Or9 – NS.

Mosquito olfactory receptors are thought to play a central role to receive and communicate the chemical message to higher brain centre through olfactory receptor neurons for decision making events (10), (42), (43). To further clarify this molecular relation, initially, we catalogued 50 different chemosensory receptors (Table-2) comprising odorant receptors (ORs); gustatory receptors (Grs) and variant ionotropic receptors (IRs), which appeared predominantly in the naïve and blood fed cohorts of the RTSS category (Table: S4). Interestingly, a cluster of 19 different olfactory receptor genes was found to be expressed abundantly and exclusively in the naïve mosquito [Table-S4]. At the same time, we also observed that a distinct repertoire of chemosensory receptor genes uniquely appeared in the blood fed cohorts, but their number is much lower than the naïve mosquito (Table S4). Observation of the constitutive expression of Orco and few other ORs and Grs (totalling 10 transcripts) in all the experimental conditions highlighted the importance of Orco for the presentation of other receptors in the olfactory system. Together, these data suggested that an abundant expression of olfactory receptors in naïve mosquitoes may be essential to encounter and manage different conflicting behavioral demands when changing from naïve sugar feed status to blood fed.

**Table 2.**

| Sl. No. | Sample Name | Number of Olfactory Receptors Transcripts |
| --- | --- | --- |
| 1. | Ac-OLF-Naive | 32 |
| 2. | Ac-OLF-30min PBM | 11 |
| 3. | Ac-OLF-30hr-PBM | 7 |
|  | Total | 50 |

Unlike OBPs, poor modulation of olfactory receptor gene expression under circadian rhythm (**Fig. 5c**) suggested a minimal role of ORs in the initialization of host-seeking behavioral activities. Alternatively, we also interpreted that ORs may not have direct biphasic regulation, but may influence a successful blood feeding event. To further corroborate with the above propositions and uncover the functional correlations of olfactory receptor responses, we monitored the transcriptional regulation of the selected OR transcripts in response to two consecutive blood meal series follow-up (44). An age-dependent enrichment of OR transcripts till 6^th^ day of maturation in the sugar-fed mosquitoes suggested that naïve mosquitoes may express and attain a full spectrum of chemosensory genes expression to meet all the needs of their life cycle requirements i.e. host seeking and mate-finding behavioral response (**Fig. 5d**).

First blood meal of 6^th^-day old naïve mosquitoes initiates the suppression of almost all the olfactory receptor transcripts within 30 minutes of blood feeding, whose expression almost ceased to a basal level at 30 hrs post blood meal, except the slight up-regulation of two transcripts named OR42 and OR62 (**Fig. 5d**). Apparently, after 30 hrs PBM we observed a significant modulation of receptor gene expression which started enriching till 72 hrs of post first blood meal, a time window coincides with the successful completion of the oviposition event. However, we did not observe any significant change in the expression of the receptor transcripts in response to second blood meal (**Fig 5d**). These findings strongly suggested that first blood meal exposure to odorant receptors may have priming effect over host seeking behavioral activities, enabling mosquito for rapid blood meal uptake for consecutive gonotrophic cycles.

**Blood meal response to other olfactory proteins:** Encouragingly, the above data prompted us to test transcriptional profiling of few uncharacterized chemosensory class of olfactory proteins, identified from the transcriptomic data (**Table S5**). Transcripts encoding orphan receptor R21, scavenger receptor class B (SRCB), an uncharacterized Protein (XP_001959820) and Sensory neuron membrane protein (SNMP) showed a similar pattern of regulation, suggesting that a combination of all the receptor type represented in the olfactory tissue of *An. culicifacies* mosquito function concurrently in nature’s aroma world and changed significantly prior and after the first blood meal as compared to the consecutive second blood meal (**Fig. 6a**). The involvement of G-proteins and related metabotropic signalling mechanism in the olfactory signal transduction of insects remain controversial. However, a rapid and consistent induction of adenylyl cyclase gene after 30m PBM (**Fig. 6b**), supporting the previous hypothesis that the synthesis of the secondary messenger, cAMP by adenylate cyclase, facilitates odorant mediated signal transduction process which further influence downstream behavioral responses (10). Surprisingly, a finding of <1% of transcripts encoding putative immune proteins, suggested the maintenance of a basal level of sterility is essential for proper olfactory functions (**Fig. S6**).

**Figure 6:**
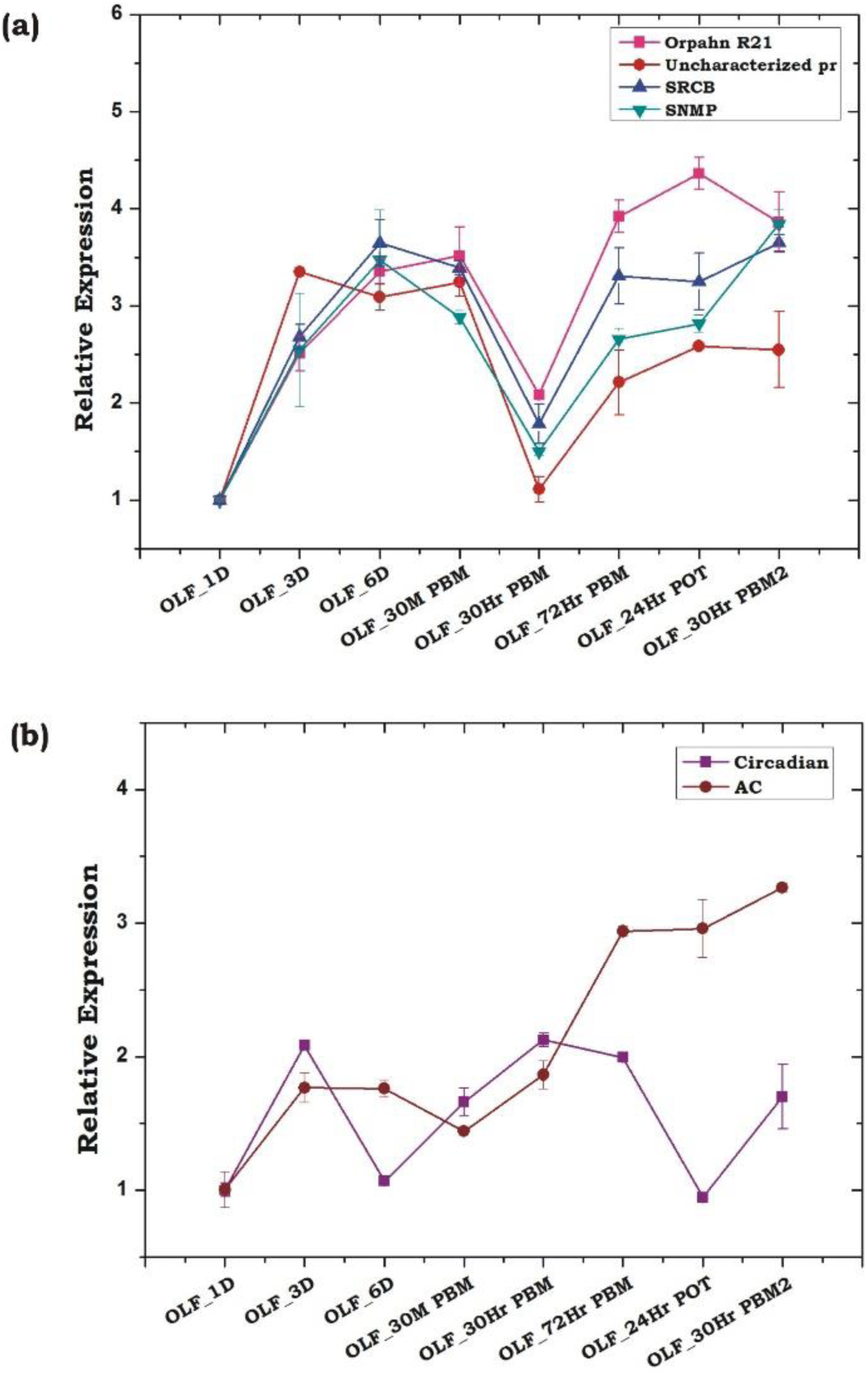
Transcriptional responses of other olfactory genes hypothesized to play crucial role in host seeking and blood feeding behavior. (a) Relative expression profiling of other receptor genes according to blood meal time series (described in fig. 5). Orphan R21: Orphan receptor 21; Uncharacterized Pr: Uncharacterized protein; SRCB: Scavenger Receptor class B; SNMP: Sensory neuron membrane protein. (b) Transcriptional profiling of other signalling molecule in response to blood meal time series experiment. Circadian: Circadian gene; AC: Adenylyl cyclase. The significance of suppression of other olfactory genes expression after 30hrs of post blood meal are as follows: Orphan R21 – ≤ 0.002; Uncharacterized pr – ≤ 0.002; SRCB – ≤ 0.006; SNMP – ≤ 0.007.

**Sensory appendages proteins as a unique target to *Anopheles culicifacies*:** Two mosquito vector species are predominant in India viz. *An. stephensi and An. culicifacies*, the former is the urban vector and later is the rural one (28), (45). *An. culicifacies* have been reported predominantly as zoophilic in India, while *An. stephensi* exhibit predominantly anthropophilic behaviour (28), (46), (47). Even, when reared under the same environment at the central insectary facility, still they display a significant difference in their behavioral properties such as feeding, mating, biting preferences etc. (personal observation/ST-S5). Though the molecular basis of such biological variation is yet to unravel, but emerging evidence suggest a significant genetic difference exists among various Anopheline mosquito species, including *An. stephensi* and *An. culicifacies* (48), (49). Under laboratory investigation, we frequently observed that biological rhythm may have a significant influence on the biting and blood feeding behavior of *An. culicifacies*. Previously, several odorant binding proteins such as OBP20/OBP1/OBP7 have been characterized as a key molecular target in many *Anopheline* mosquitoes involved in host-seeking behavior (50), (51), (52), but remains unidentified from *An. culicifacies*.

Thus, to test whether any species-specific olfactory derived genetic factors have any differential regulation, influencing the behavioral responses, we compared the expression of at least 6 OBPs transcripts between two laboratories reared mosquito species *An. stephensi* and *An. culicifacies*. In this analysis we also included two novel SAP proteins, which showed a high induction than other OBPs in the olfactory system of the mosquito *An. culicifacies* at midnight (**Fig. 4a**). Surprisingly, a sex and tissue specific comparative transcriptional profiling of selected OBPs revealed a dominant expression of SAP1 (p *≤* 0.0003)/SAP2 (p *≤* 0.0007) in the legs of mosquito *An. culicifacies* (**Fig 7a, b**). Together these data indicated that *An. culicifacies* may draw an extra advantage of having more sensitive appendages, possibly to favour more active late night foraging behavior, than *An. stephensi*. Furthermore, a strong phylogenetic association of Ac-SAP1 protein within Anthropophilic mosquito clade (**Fig. 7c**), strongly suggested that sensory appendages proteins may have a crucial role to meet and manage the high host seeking behavioral activities, restricted to *An. culicifacies*.

**Figure 7:**
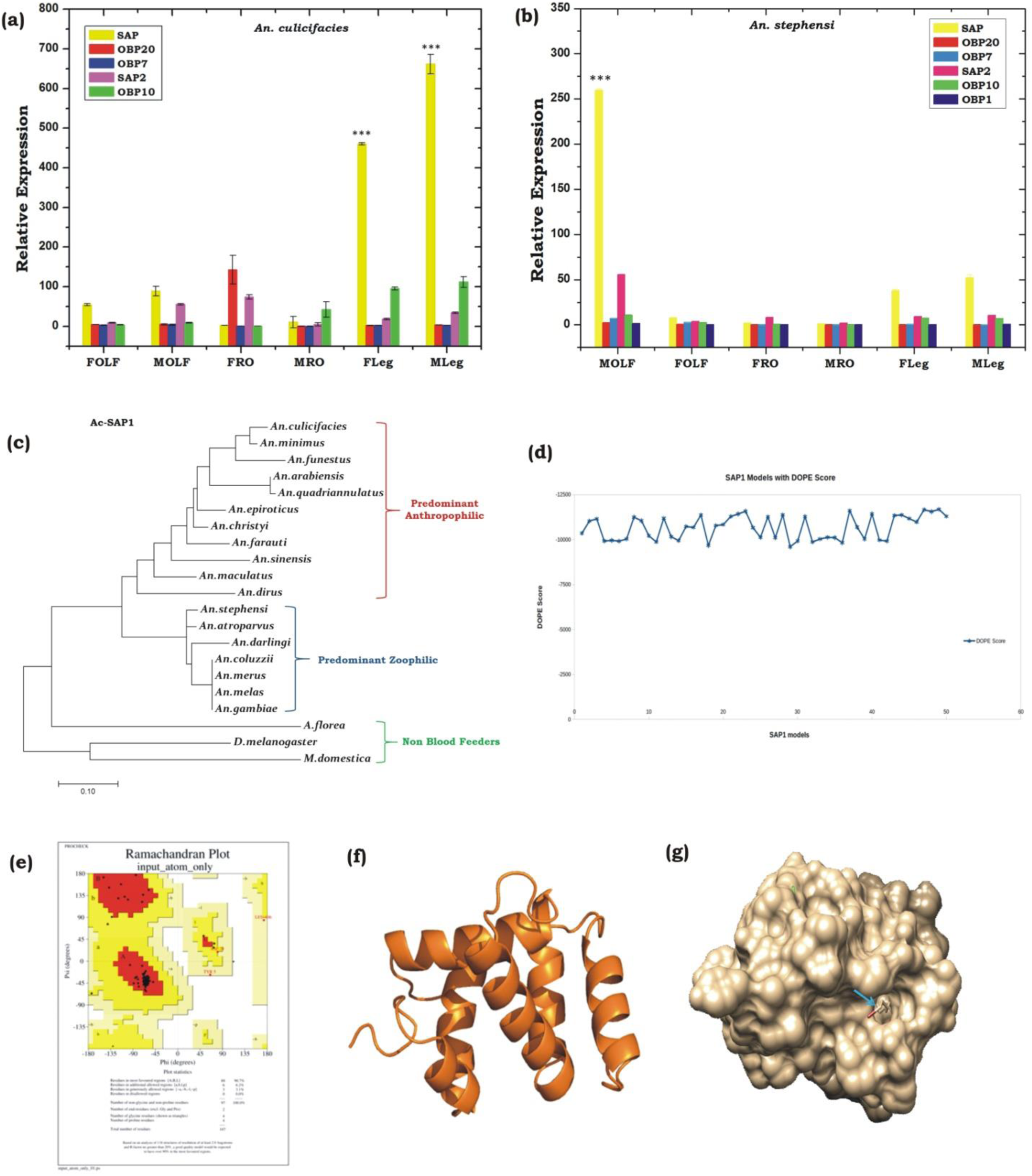
Comparative transcriptional responses of Odorant binding protein genes between two major Indian vectors and structural characterization of one of the potent OBP gene. (a, b) Sex and tissue specific relative expression profiling of OBP genes in *An. culicifacies* (a) and *An. stephensi* (b). FOLF: female olfactory tissue (OLF); MOLF: Male OLF; FRO: Female reproductive organ; MRO: Male reproductive organ; FLeg: Female legs; MLeg: Male legs. OBP gene details: SAP: Sensory appendages protein 1; SAP2: Sensory appendages protein 2. (c) Phylogenetic analysis of *An. culicifcaies* SAP1 (Ac-SAP1) gene. (d) DOPE score analysis for SAP1. (e) Ramachandran Plot of SAP1 protein. (f) 3-dimentional protein structure of Ac-SAP1 protein. (g) Binding site of SAP1 protein showed in space fill with nearby residues in stick form.

Encouraging to the above finding, we carried out a 3D structure modelling analysis of Ac-SAP1 and Ac-SAP2, to predict the best possible conserved binding pockets for specific chemicals. In the absence of any available solved X-ray structure of reference SAP protein, we applied a template based comparative molecular modelling approach. An initial blast analysis identified two best templates in PDB database code for chemosensory protein 2GVS and 1KX8 with identity 47-56% and coverage >80%, favouring their suitability for structure prediction. Out of the 50 modelled 3D structures for each protein, DOPE score analysis resulted in the selection of model-49 and model-27 with score −11689.73, and −10989.75 for SAP1 and SAP2 respectively (**Fig. 7d and Fig. S7a**).

We validated the best-selected model using Procheck server for Ramachandran plot, showing a more than 95% allowable region, with no residue falling in the disallowed region of the plot (**Fig. 7e and Fig. S7b**). Based on the consensus, a best-fit ligand binding site prediction analysis within the selected models was scored by COACH server, which engages at least five different algorithms TM-SITE, S-SITE, COFACTOR, FIND-SITE, and ConCavity. Binding pocket for SAP1 and SAP2 identified eight consensus residues namely D36, E39, L40, K49, C52, Q59, Y91, and Y95 along with BDD (12-bromo-1-dodecanol) as a predicted ligand. Minimization of the steric clashes from the complex structures was done using Chimera software (**Fig. 7f and Fig. S7c**). Furthermore, selection of amino acid residues within 3 Å region of ligand molecule are I43 and Y95 of which Y95 is involved in H-bonding with BDD ligand. Similarly, in case of SAP2 protein, residue selection resulted in the identification of I43, D51, Q59, T63, Y95 residues of which D51 form H-bond with BDD ligand (**Fig. 7g and Fig. S7d**).

In our analysis we observed the presence of at least two conserved cysteine (CYS52 and CYS55) residues in the loop region of SAP1 and SAP2 proteins, which may likely involved in di-sulphide bond formation and stabilization of protein structure. Our analysis also showed that binding pocket forms a tunnel-like structure which is preferred by long aliphatic molecules. Presence of negatively charged asparatic and glutamic acid at both ends showed the preference for charged residue near the vicinity of ligand molecule. Moreover, the presence of conserved negatively charged asparatic acid and polar tyrosine (TYR91 in SAP1 and TYR95 in SAP2) at one end of binding pocket suggested their role in ligand binding.

**A synergetic action of OBPs/ORs may manage behavioral responses:** Decoding the genetic relationship of sense of smell is central to design new molecular tools to disrupt mosquito-human interaction. We demonstrated that a synergistic and harmonious action of olfactory encoded unique factors govern the successful ‘prior and post’ blood feeding associated behavioral complexities. We concluded that a quick recovery of the actions of odorant binding proteins immediately after blood feeding, and delayed re-activation of olfactory receptor proteins after blood meal digestion completion are unique to manage diverse behavioral responses. We hypothesize that first blood meal exposure is enough for prime learning, satisfying the motivational search of mosquitoes for the completion of their gonotrophic cycles. Thus, it is plausible to propose that apart from the innate odor responses, adult female mosquitoes might took an advantage of prior odor (vertebrate) exposure, which leads an exclusive evolutionary specialty, allowing them to learn, experience and adapt as a fast blood feeder in nature (Fig. 8).

**Figure 8:**
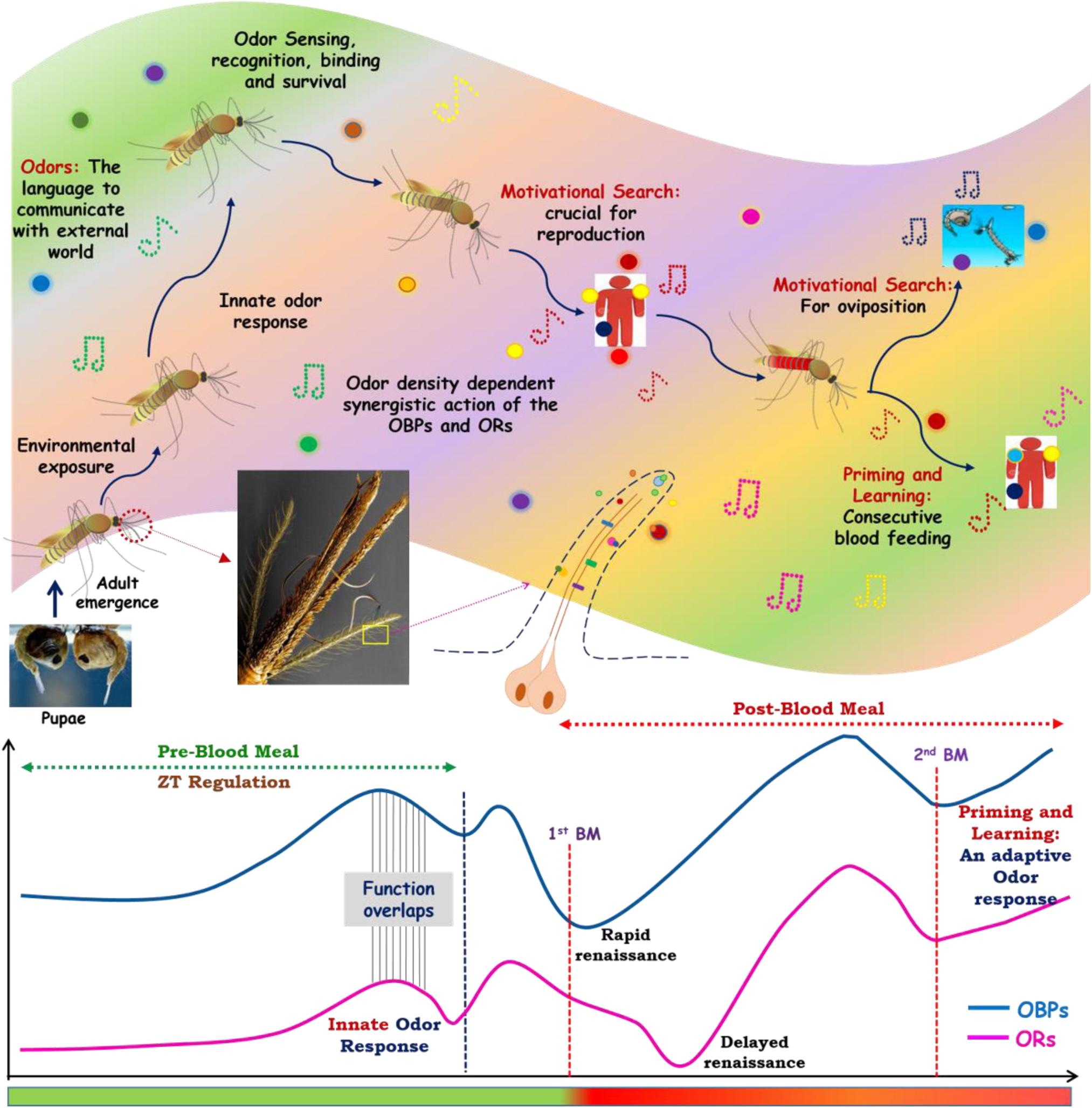
How smart actions of olfactory system manages blood feeding associated odor response: an evolutionary speciality of adult female mosquitoes. After emergence from pupae adult mosquitoes are exposed to the overwhelmed odor world, where odorants chemicals act as a language of communication with the external world. The sophisticated innate olfactory system of mosquitoes enables them to recognize and differentiate these wide variety of odorants which are crucial for their every life cycle stages. Inner physiological motivation as well as the age and exposure of mosquitoes towards the external world promote them for host seeking and blood feeding event. After taking blood meal mosquitoes initiate next level of physiological cum behavioral events i.e. oviposition. Apart from that, first exposure with vertebrates facilitate learning and second blood feeding events. These whole odors mediated response is tactfully managed by the synergistic actions of Odorant binding proteins (OBPs) and olfactory receptors (Ors). The overlapping circadian rhythm dependent functions of OBPs and Ors govern the pre blood meal events of host fetching events. As soon as the mosquitoes take blood meal the functions of OBPs and Ors ceased for some period, but the recovery of OBPs actions occurs early as compared to Ors to perform the next level of behaviors. Mosquitoes, then take advantage/adapted from priming and learning of the first blood meal exposure for the more rapid consecutive blood feeding.

In summary, we decoded and established a possible functional correlation that how a coherent and smart actions of olfactory encoded factors enabled adult female mosquitoes to meet and manage the blood feeding associated complex behavioral activities (**Fig. 8**). Furthermore, targeting species-specific unique genes such as sensory appendages proteins may be crucial to design disorientation strategy against mosquito *An. culicifacies*, an important malarial vector in rural India.

## Funding statement

Laboratory work was supported by Indian Council of Medical Research (ICMR), Government of India (No.3/1/3/ICRMR-VFS/HRD/2/2016) and Tata Education and Development Trust (Health-NIMR-2017-01-03/AP/db). Tanwee Das De is the recipient of UGC Research Fellowship (CSIR-UGC-JRF/20-06/2010/(i) EU-IV. The funders had no role in study design, data collection and analysis, decision to publish, or preparation of the manuscript.

## Data Deposition

The sequencing data were deposited to National Center for Biotechnology Information (NCBI) Sequence Reads Archive (SRA) system (BioProject accessions: <u>PRJNA414162</u>; BioSample accessions: SAMN07981002, SAMN07972755, and SAMN07775994).

## Authors contribution statement

Conceived and designed the experiments: TDD, RD; Performed the Experiments: TTD, TT, SV, DS, VS, PS, CR, SK, ST, JR; Analyzed the data: TDD, RD; YH, Contributed reagents/materials/analysis tools: YH, RD, KCP; Wrote the paper: TDD, RD, YH, KCP.

## Competing interest statement

The authors declare no conflict of interest

## Acknowledgement

We thank insectary staff members for mosquito rearing. We also thank Kunwarjeet Singh for technical assistance in laboratory. Finally, we are thankful to Xceleris Genomics, Ahmedabad, India for generating NGS sequencing data.

## Supporting Information

**Fig. 1.**
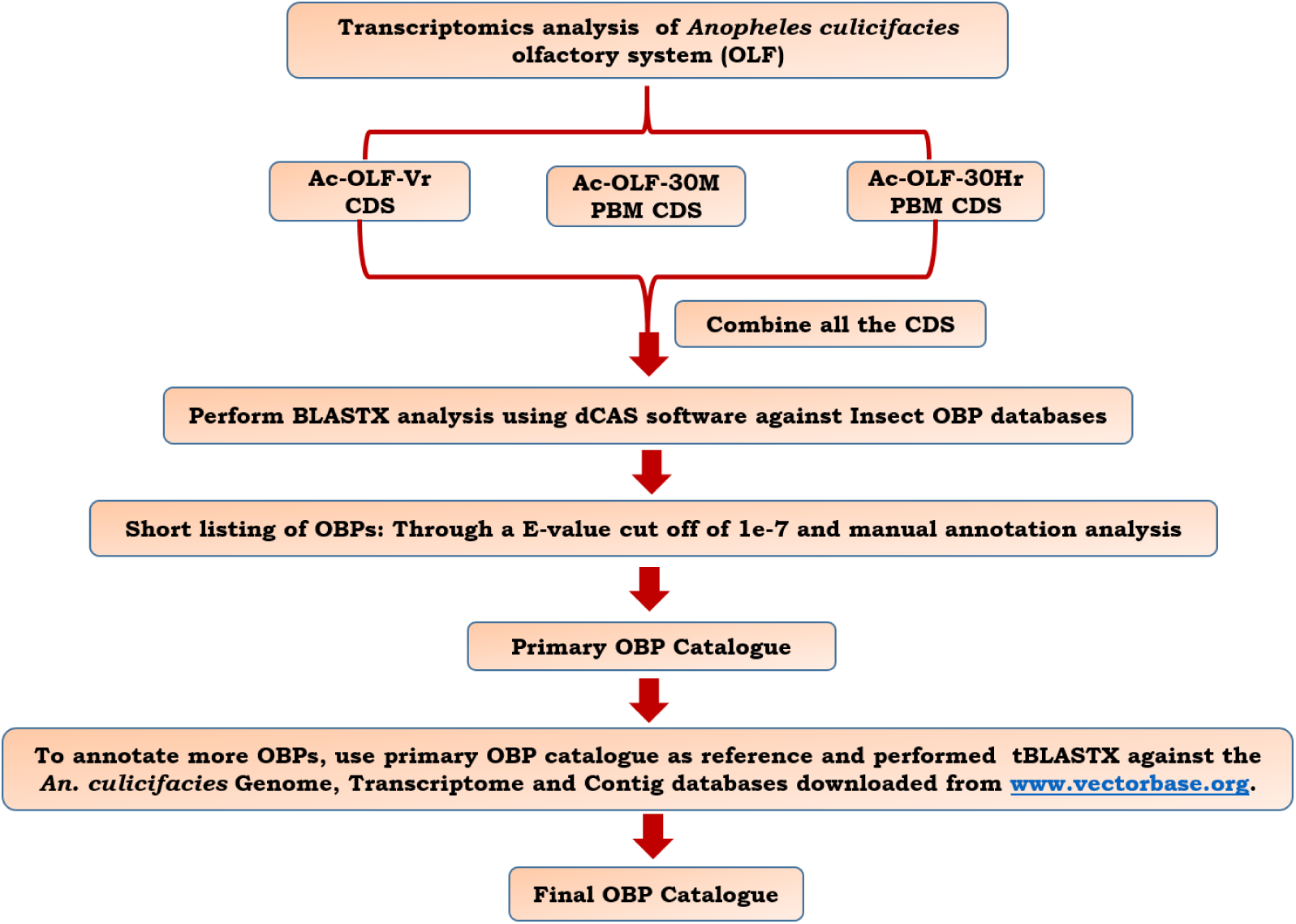
Bioinformatic workflow followed to identify the odorant binding protein (OBPs) genes of *Anopheles culicifacies* mosquito. Putative OBP transcripts were identified from olfactory tissue transcriptome database by performing BLASTX analysis against the insect OBP database using dCAS software package with an E-value cutoff 1e^−7^. To identify and annotate more putative OBP genes from genome of *An. culicifacies*, we performed tBLASTX analysis of all shortlisted OBPs as a query against genome database, downloaded from <u>www.vectorbase.org</u>. Removal of all kinds of repetition was done and prepare the final OBP catalogue.

**Fig. S2:**
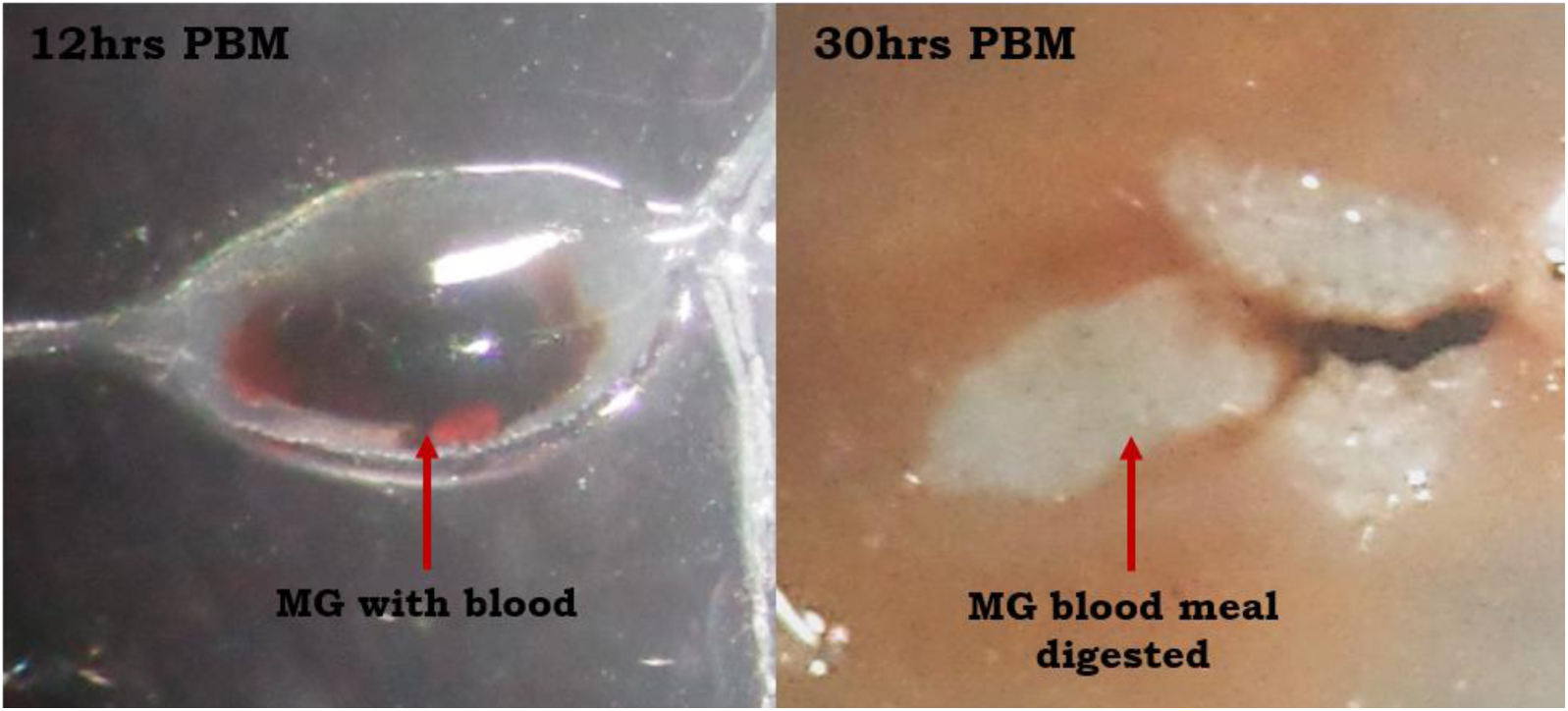
Time dependent depiction of blood fed midgut (MG) showing blood meal digestion. MG digested from 12hrs post blood meal (PBM) showed undigested blood still present in the MG but after 30hrs of PBM blood meal digestion completed and thus no blood remnants is observable.

**Fig. S3:**
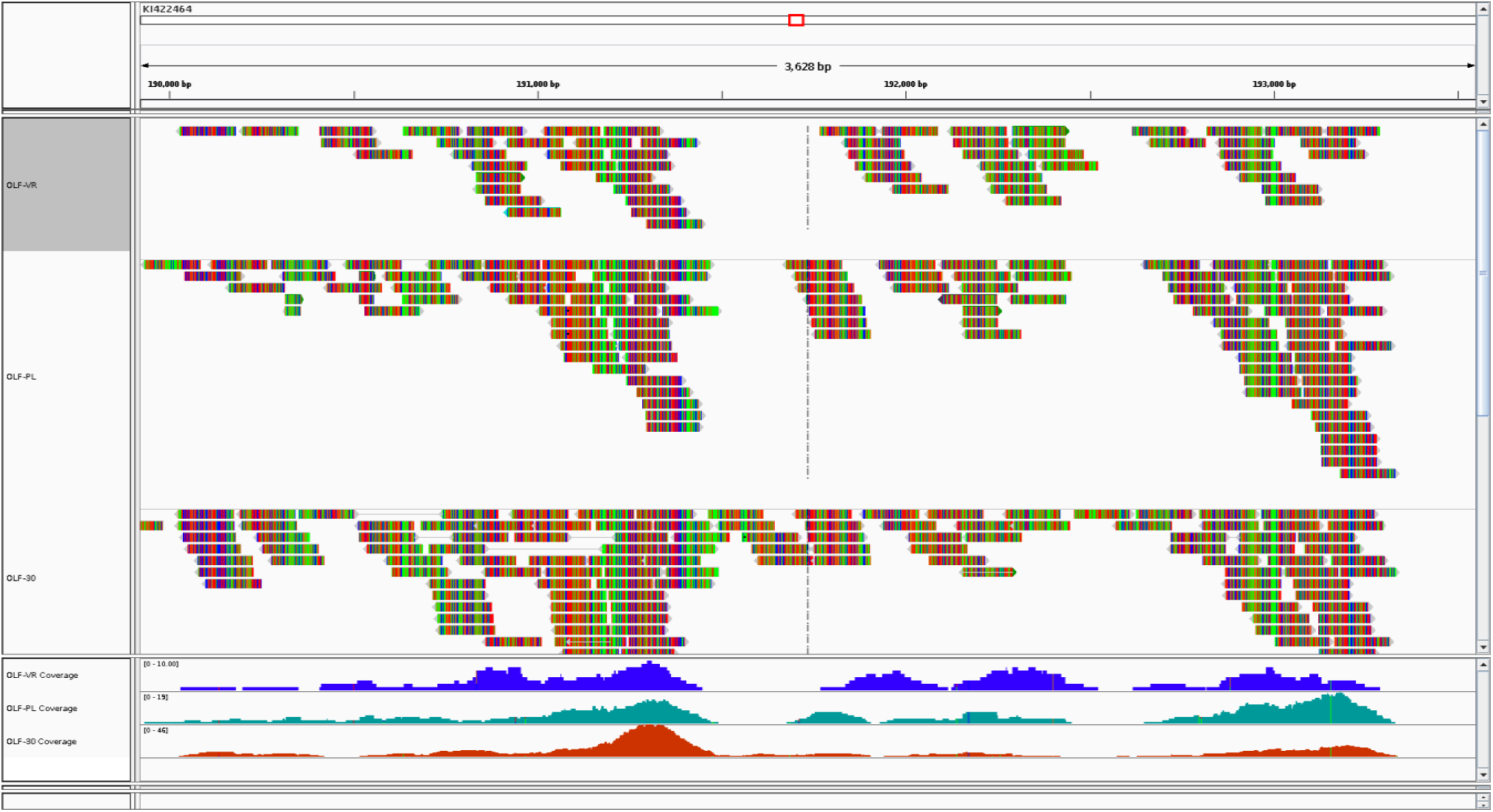
Self-explanatory pictorial presentation of genome guided reference mapping limitations of olfactory database of *Anopheles culicifacies*.

**Fig. S4a:**
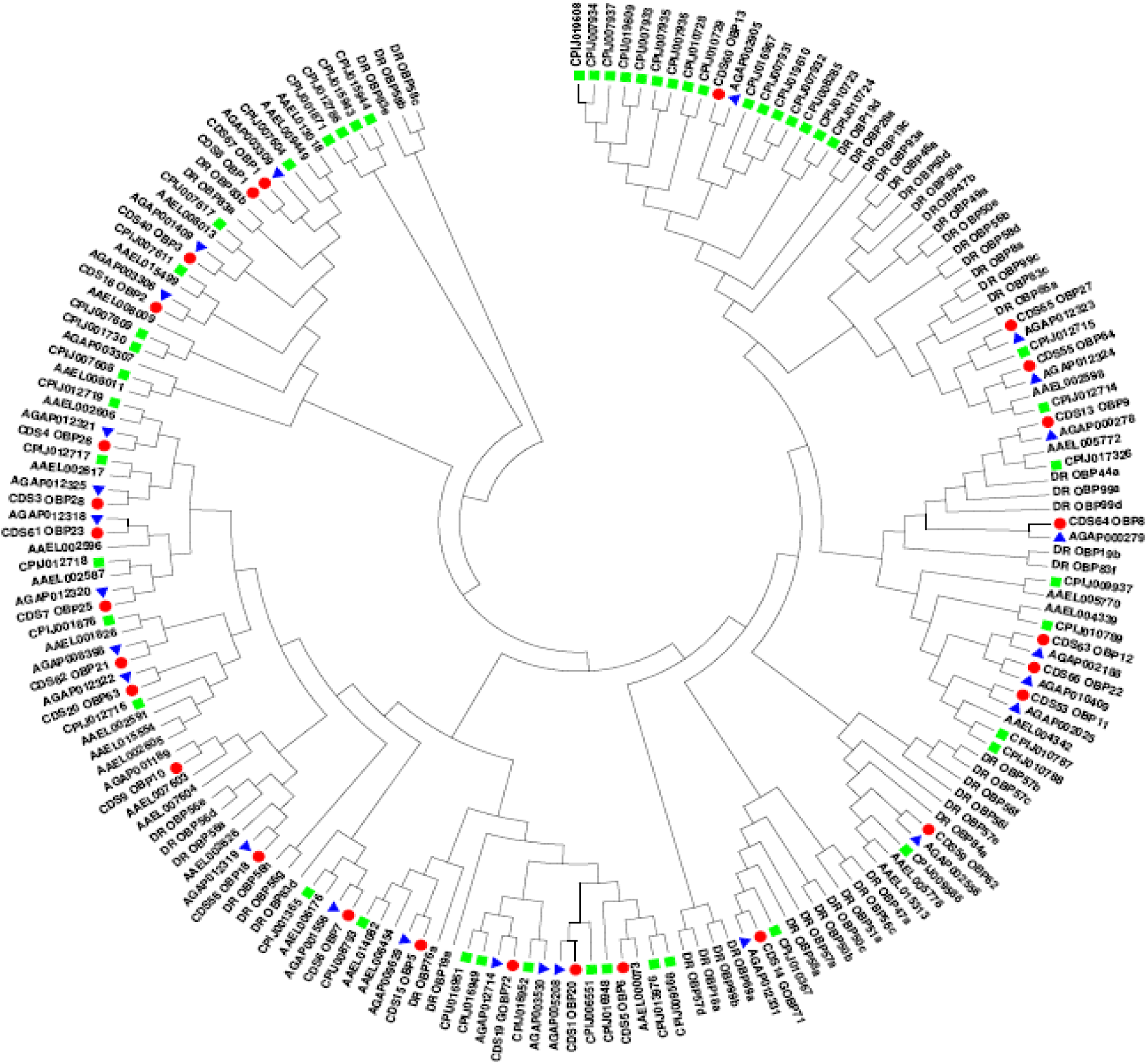
Phylogenetic analysis of Classic-OBPs. Classic-OBPs of *An. culicifacies* showed conserved sequence relationship with *An. gambiae* and other mosquito and insect species. Different color code that indicating a particular mosquito and insect species: Blue – *An. gambiae*; Red – *An. culicifacies*, Green – *Culex quinquefasciatus*. *Drosophila melanogaster* (DRX) and *Aedes aegypti* (AAX) are not marked with any color code.

**Fig. S4b:**
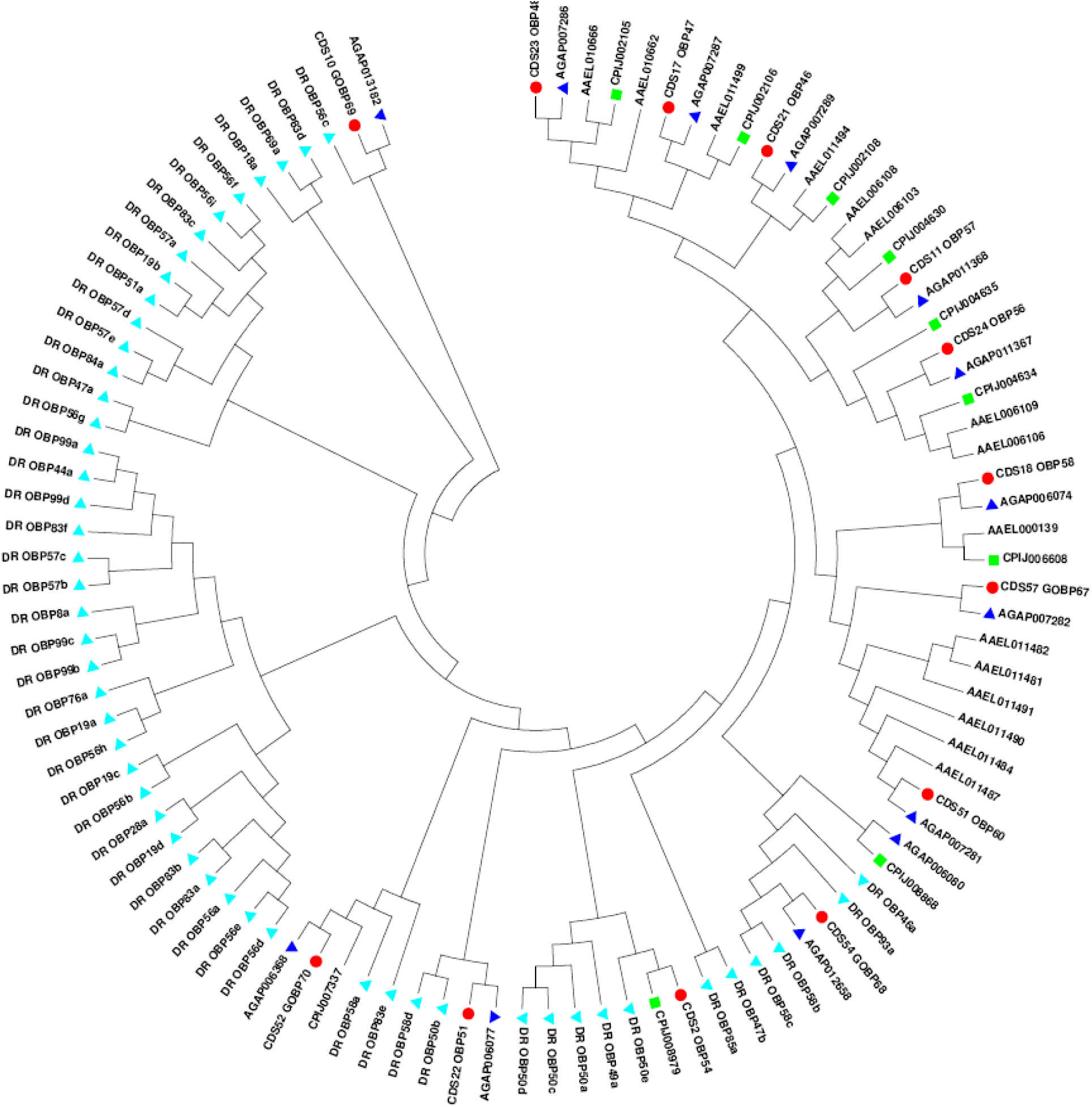
Phylogenetic analysis of Plus-C OBP superfamily. *An. culicifacies* Plus-C class of OBPs are more closely linked to mosquito specific OBPs and showed distant relationship to other non-mosquito species for example *Drosophila.* Different color code that indicating a particular mosquito and insect species: Blue – *An. gambiae*; Red – *An. culicifacies*, Green – *Culex quinquefasciatus*; Sky blue – *D. melanogaster. Aedes aegypti* (AAX) is not marked with any color code.

**Fig. S4c:**
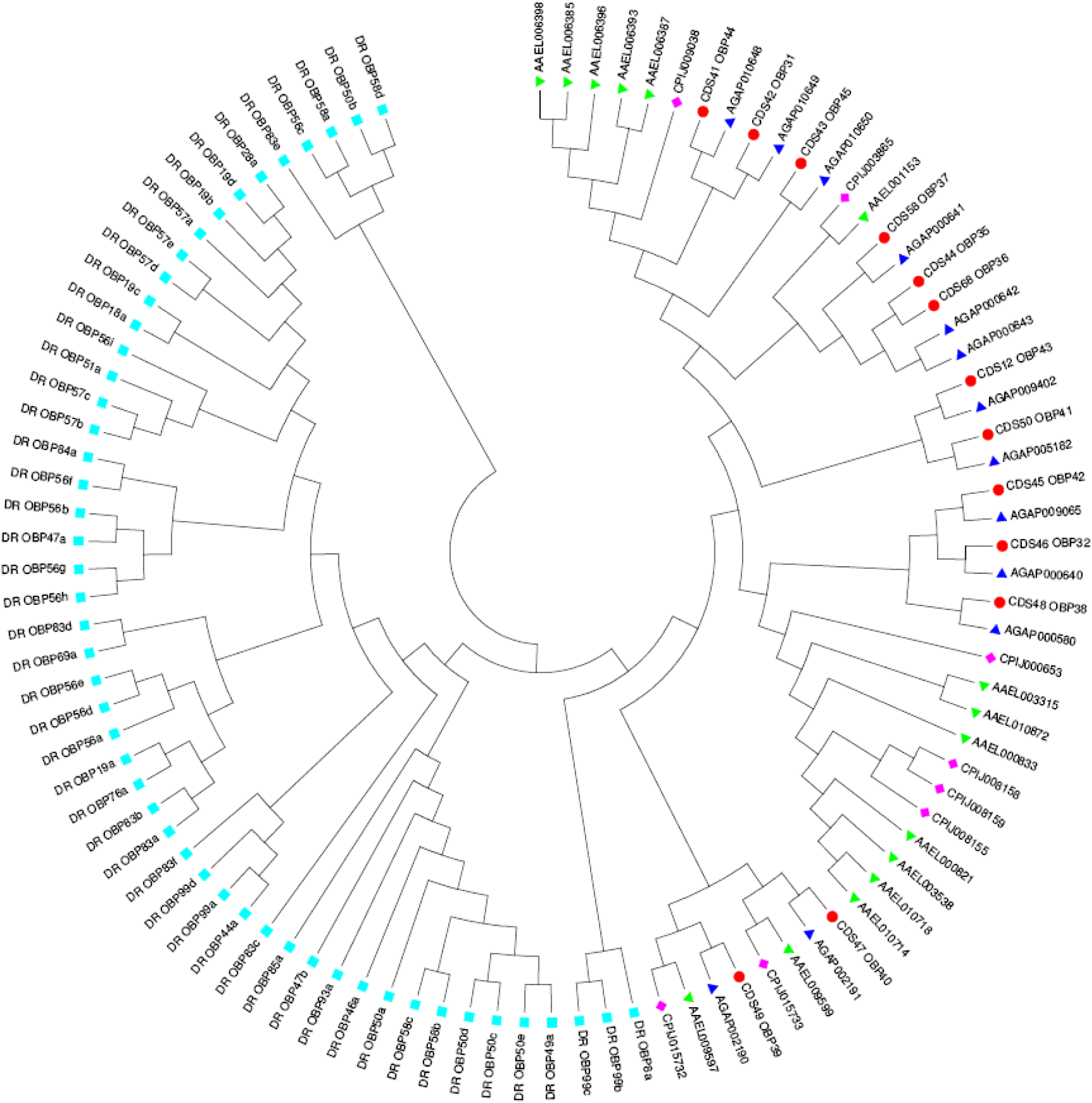
Phylogenetic analysis of Atypical OBP superfamily. *An. culicifacies* Atypical OBP class are more closely linked to different mosquito species and showed distant relationship to other non-mosquito species for example *Drosophila.* Different color code that indicating a particular mosquito and insect species: Blue – *An. gambiae*; Red – *An. culicifacies*, Pink – *Culex quinquefasciatus*; Sky blue – *D. melanogaster;* Green - *Aedes aegypti*.

**Fig. S5:**
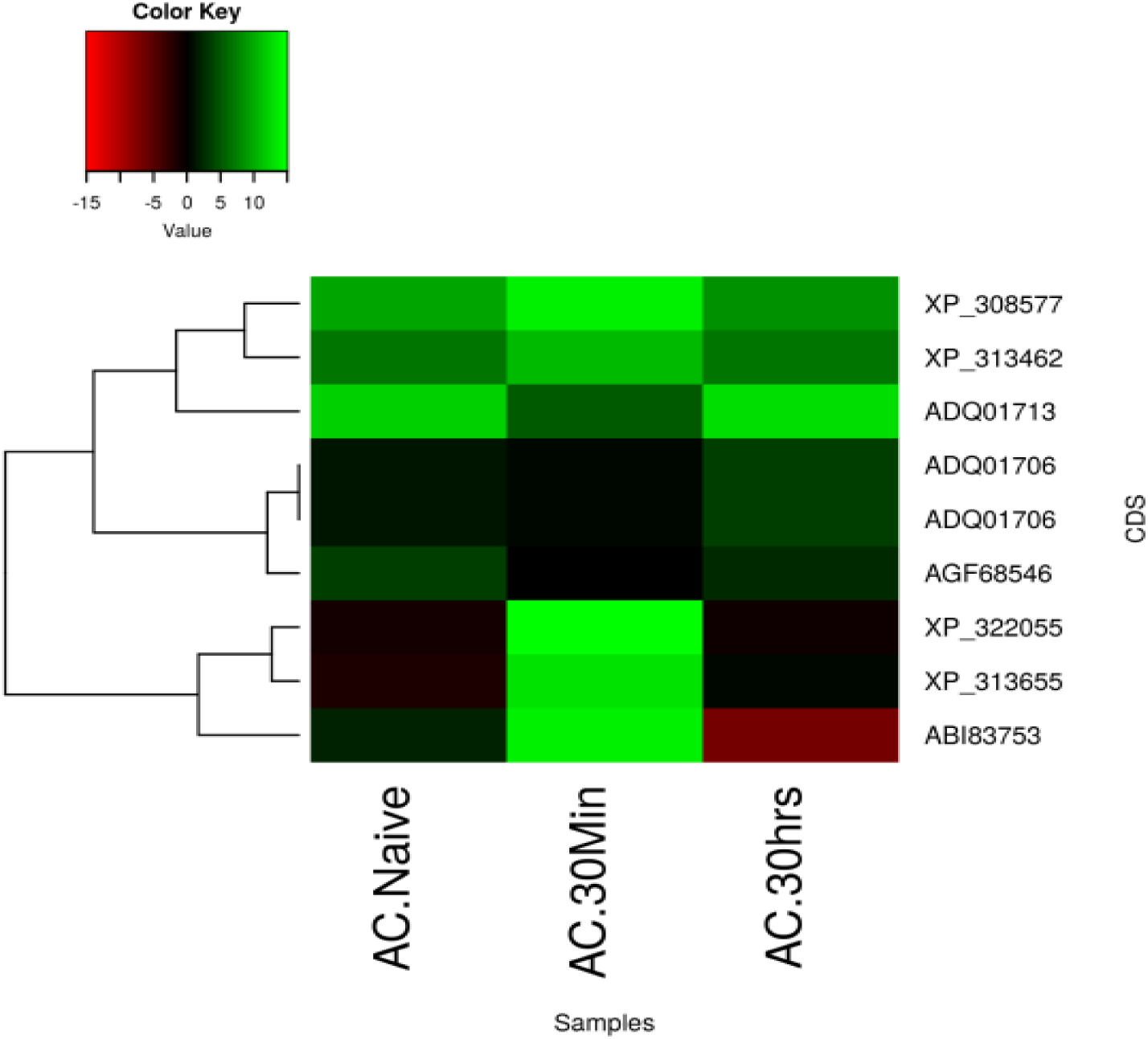
Transcriptional Response of *An. culicifacies* OBPs. (a) Heat map showing differential expression pattern of eight common OBP genes in naïve and blood fed olfactory tissue of *An. culicifacies*.

**Fig. S6:**
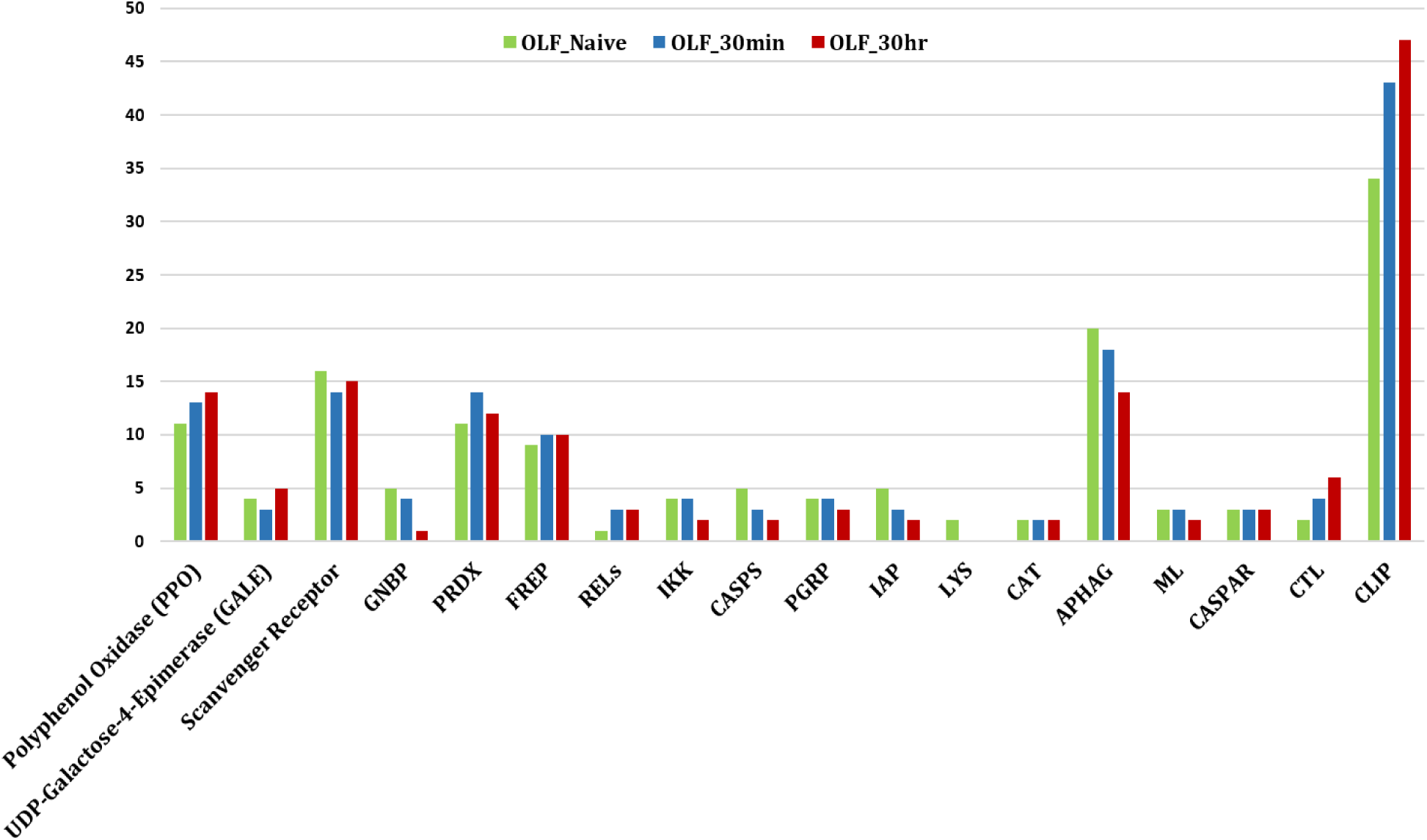
Differential expression pattern of olfactory specific immunome. The immune genes expressed in the olfactory system of *An. culicifacies* are retrieved through BLASTX analysis against the insect ImmunoDB database and categorized into eighteen different family members and their differential expression pattern was determined by the number of sequences appeared in each RNASeq data of naïve and blood fed mosquitoes.

**Fig. S7:**
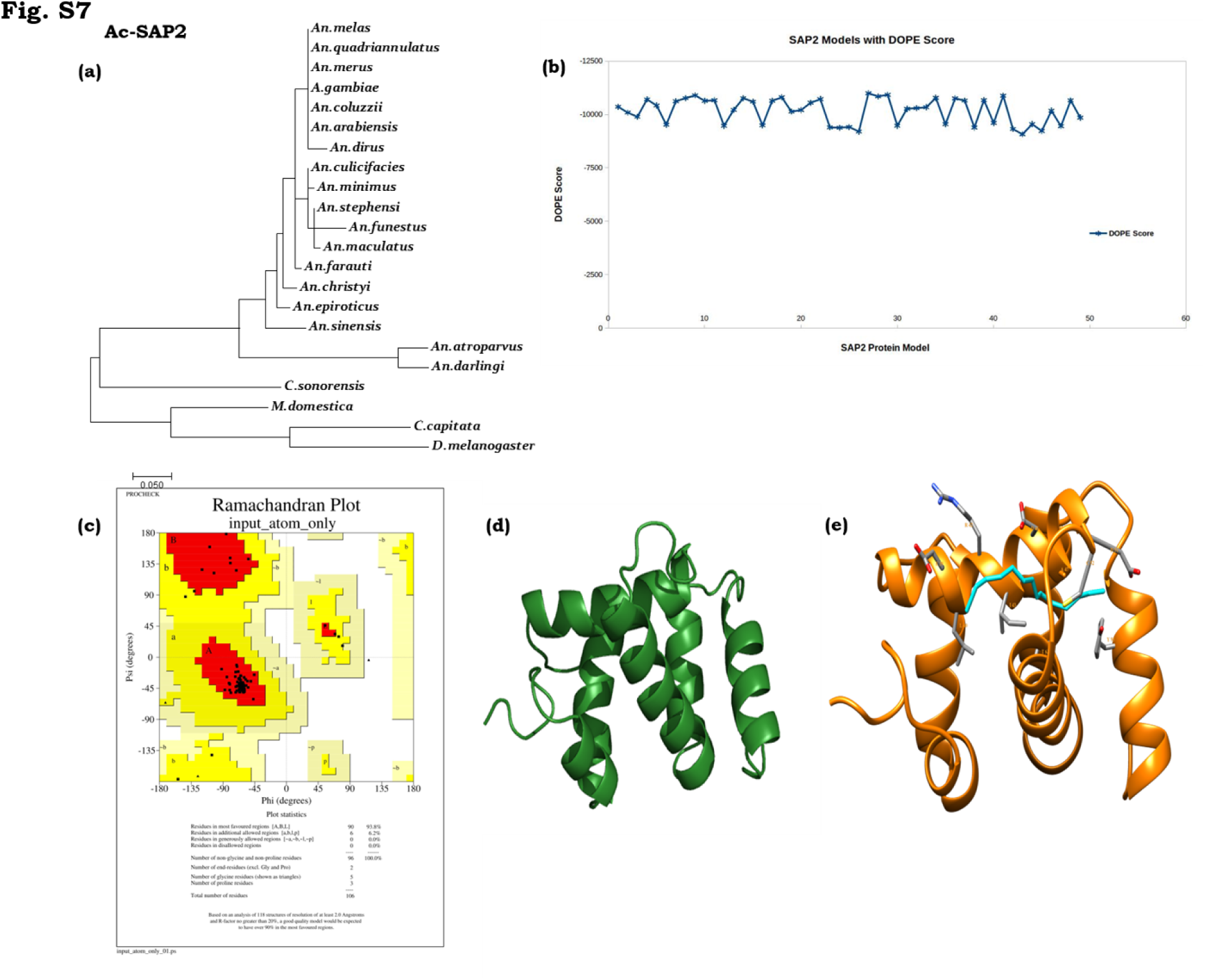
Phylogenetic and structural analysis of SAP2. (a)Phylogenetic analysis of *An. culicifcaies* SAP2 gene (Ac-SAP2). (b) DOPE score analysis for SAP2. (c) Ramachandran Plot of SAP2 protein. (d) 3-dimentional protein structure of Ac-SAP2 protein. (e) Binding site of SAP2 protein showed in space fill with nearby residues in stick form.

**Supplementary Table 1:** Annotation kinetics of the RNA-Seq data.

| **Molecular Features** | **Ac-OLF-Naive** | **Ac-OLF-30M PBM** | **Ac-OLF-30Hr PBM** |
| --- | --- | --- | --- |
| **Total Transcripts** | 8133 | 8907 | 8396 |
| **Total BLASTx hits (NR)** | 7245 (~89%) | 7553 (~84.79%) | 6829 (~81.33%) |
| **Transcripts with GO Match** |  |  |  |
| **Molecular Function** | 3946 | 4137 | 3763 |
| **Biological process** | 3804 | 3985 | 3566 |
| **Cellular component** | 2097 | 2201 | 1999 |
| **Transcript with KEGG match** | 2645 (~32.52%) | 2975 (~33.40%) | 2740 (~32.63%) |

**Supplementary Table 2:**
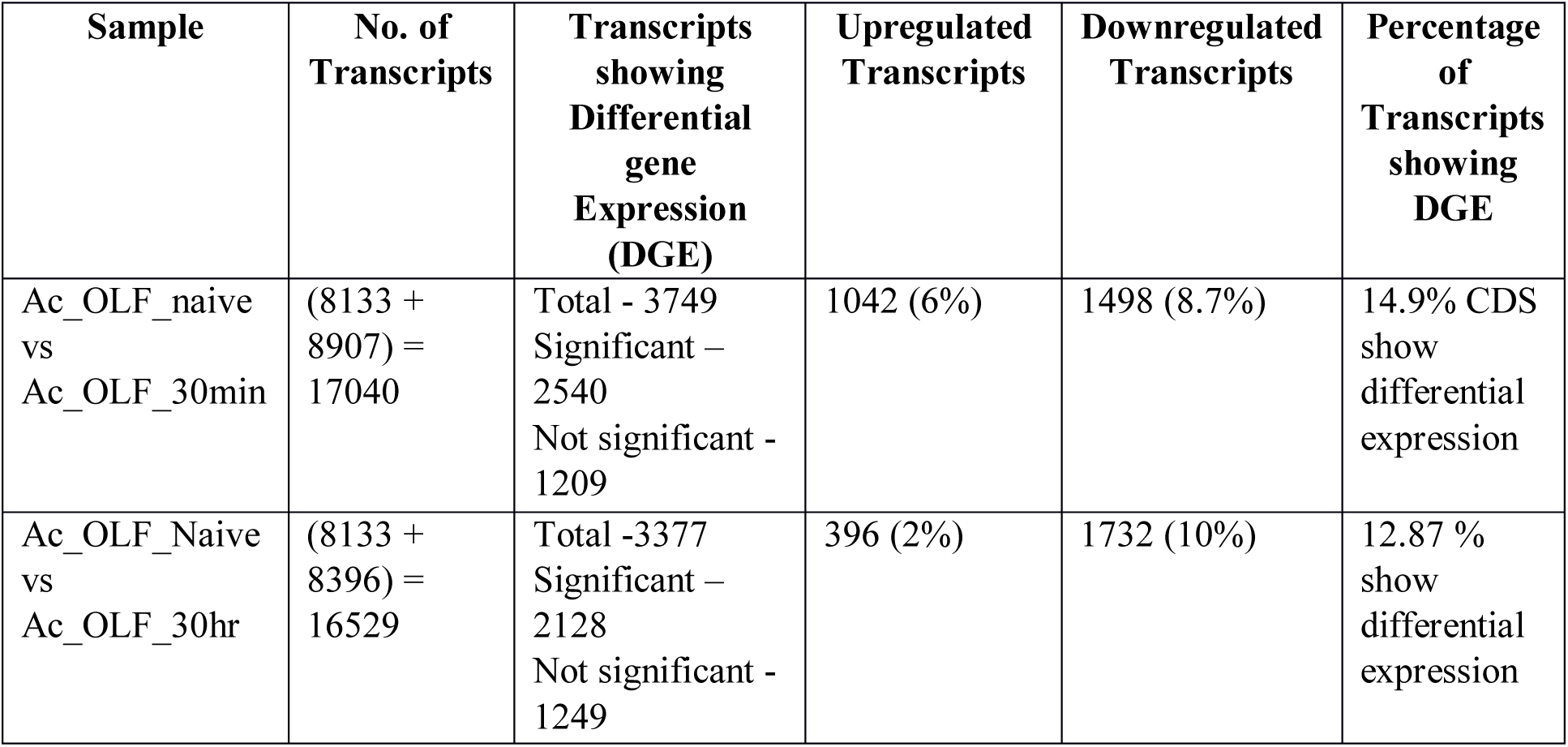
Percentage of Differentially Expressed Transcripts.

**Supplementary Table S3:** Genome Annotated OBP.

| Serial No. | Transcript ID | Origin of the Transcript | Name | *An.culicifacies* Database Match | E-Value | Identity | Sub-Family | Ortholog in *An.gambiae* | FPKM |
| --- | --- | --- | --- | --- | --- | --- | --- | --- | --- |
| 1. | Transcript _1018 | 30hr PBM | OBP20 | ACUA000236 | 7E-63 | 95.70 % | Classic | AGAP005208 | 3173.27822520792 |
| 2. | Transcript _1216 | 30hr PBM | GOBP72 | ACUA019958 * | 6.00E-57 | 99% | Classic | AGAP012714 | 1254.82586 |
| 3. | Transcript _127 | 30hr PBM | OBP7 | ACUA012847 | 2.00E-85 | 98% | Classic | AGAP001556 | 4144.93009 |
| 4. | Transcript _1450 | 30hr PBM | OBP26 | ACUA003587 * | 3.00E-80 | 98.50 % | Classic | AGAP012321 | 2051.81759 |
| 5. | Transcript _4885 | 30hr PBM | OBP63 | ACUA020502 * | 6.00E-70 | 99.30 % | Classic | AGAP012322 | 187.690564 |
| 6. | Transcript _6613 | 30hr PBM | OBP3 | ACUA024106 | 5.00E-07 | 92.30 % | Classic | AGAP001409 | 21144.6696 |
| 7. | Transcript _91 | 30hr PBM | GOBP71 | ACUA003567 | 9.00E-169 | 95.50 % | Classic | AGAP012331 | 4939.85295 |
| 8. | Transcript _109 | Naive | OBP5 | ACUA003763 | 2.00E-48 | 97.60 % | Classic | AGAP009629 | 301.156325 |
| 9. | Transcript _3113 | Naive | OBP28 | ACUA001691 | 1.00E-75 | 96.20 % | Classic | AGAP012325 | 63.5528511 |
| 10 | Transcript _1290 | Naive | OBP9 | ACUA003296 | 6.00E-89 | 100 % | Classic | AGAP000278 | 389.876797 |
| 11 | Transcript _138 | Naive | OBP6 | ACUA004501 * | 3.00E-12 | 41.20 % | Classic | AGAP003530 | 238.355131 |
| 12 | Transcript _270 | Naive | OBP2 | ACUA004189 | 8.00E-90 | 97.30 % | Classic | AGAP003306 | 211.717742 |
| 13 | Transcript _218 | 30min_PBM | OBP10 | ACUA017068 | 1.00E-71 | 99.20 % | Classic | AGAP001189 | 40.2405414 |
| 14 | Transcript _48 | 30min_PBM | OBP25 | ACUA014063 | 3.00E-65 | 99.20 % | Classic | AGAP012320 | 92.3071105 |
| 15 | Transcript _62 | 30min_PBM | OBP1 | ACUA014299 | 2.00E-87 | 99.30 % | Classic | AGAP003309 | 608.296339 |
| 16 | Transcript<br>_59 | 30min_<br>PBM | OBP54 | ACUA00<br>1499 * | 2.00E<br>-56 | 97.10<br>% | Pluc-C | AGAP00<br>6080 | 41.4381766 |
| 17 | Transcript<br>_3357 | 30min_<br>PBM | GOBP<br>69 | ACUA02<br>4897 | 3.00E<br>-94 | 93% | Pluc-C | AGAP01<br>3182 | 323.549283 |
| 18 | Transcript<br>_207 | 30min_<br>PBM | OBP47 | ACUA01<br>1874 | 2.00E<br>-115 | 96.90<br>% | Pluc-C | AGAP00<br>7287 | 116.931 |
| 19 | Transcript<br>_7419 | 30min_<br>PBM | OBP58 | ACUA01<br>9212 * | 1.00E<br>-109 | 91.60<br>% | Pluc-C | AGAP00<br>6074 | 28.093679 |
| 20 | Transcript<br>_5393 | 30min_<br>PBM | OBP46 | ACUA02<br>0810 | 8.00E<br>-123 | 95.60<br>% | Pluc-C | AGAP00<br>7289 | 23.4694097 |
| 21 | Transcript<br>_6620 | 30min_<br>PBM | OBP56 | ACUA02<br>5325 | 1.00E<br>-143 | 97.40<br>% | Pluc-C | AGAP01<br>1367 | 20.576198 |
| 22 | Transcript<br>_611 | Naive | OBP57 | ACUA02<br>8407 | 1.00E<br>-114 | 97.60<br>% | Pluc-C | AGAP01<br>1368 | 1983.04862 |
| 23 | Transcript<br>_4488 | Naive | OBP51 | ACUA02<br>3815 * | 7.00E<br>-23 | 87.50<br>% | Pluc-C | AGAP00<br>6077 | 65432.854 |
| 24 | Transcript<br>_22 | Naive | OBP48 | ACUA02<br>5211 | 3.00E<br>-59 | 87.30<br>% | Pluc-C | AGAP00<br>7286 | 1728.56394 |
| 25 | Transcript<br>_853 | 30hr<br>PBM | OBP43 | ACUA00<br>1907 | 0.00E<br>+00 | 96.30<br>% | Two<br>Domain | AGAP00<br>9402 | 90.5977525 |
| 26 | Transcript<br>_2542 | 30hr<br>PBM | D7 | ACUA02<br>5281 * | 0 | 98.70<br>% | D7<br>Protein<br>Family | AGAP00<br>6278 | 462.002781 |
| 27 | Transcript<br>_2682 | 30hr<br>PBM | CSP | ACUA00<br>8309 * | 1.00E<br>-66 | 100<br>% | Chemose<br>nsory<br>Protein | AGAP00<br>1303 | 215.86399 |
| 28 | Transcript<br>_5645 | 30hr<br>PBM | LD7 | ACUA00<br>2223 | 3.00E<br>-77 | 98.40<br>% | D7<br>Protein<br>Family | AGAP00<br>7286 | 34824.3385 |
| 29 | Transcript<br>_64 | 30hr<br>PBM | SD7 | ACUA00<br>1093 | 1.00E<br>-33 | 98.40<br>% | D7<br>Protein<br>Family | AGAP00<br>8281 | 1348.33451 |
| 30 | Transcript<br>_1808 | Naive | CSP | ACUA01<br>0232 * | 6.00E<br>-51 | 96.70<br>% | Chemose<br>nsory<br>Protein | AGAP00<br>8055 | 519.825389 |
| 31 | Transcript<br>_447 | Naive | CSP3 | ACUA00<br>4458 | 7.00E<br>-71 | 96.70<br>% | Chemose<br>nsory<br>Protein | AGAP00<br>8059 | 195.413717 |
| 32 | Transcript<br>_702 | Naive | CSP | ACUA01<br>7151 | 1.00E<br>-29 | 90.90<br>% | Chemose<br>nsory<br>Protein | AGAP00<br>8054 | 203.377554 |
| 33 | Transcript<br>_12 | Naive | Sensor<br>y<br>Appen<br>dage<br>Protein | ACUA01<br>8714 | 2.00E<br>-40 | 100<br>% | Chemose<br>nsory<br>Protein | AGAP00<br>8051 | 24346.8834 |
| 34 | Transcript<br>_5005 | Naive | LD7 | ACUA01<br>8525 | 9.00E<br>-100 | 97.70<br>% | D7<br>Protein<br>Family | AGAP00<br>8279 | 53.6723049 |
| 35 | Transcript<br>_631 | Naive | LD7 | ACUA00<br>8578 | 1.00E<br>-73 | 89.70<br>% | D7<br>Protein<br>Family | AGAP02<br>8120 | 27.625451 |
| 36 | Transcript<br>_1446 | 30min_<br>PBM | CSP | ACUA01<br>3563 * | 5.00E<br>-80 | 99.40<br>% | Chemose<br>nsory<br>Protein | AGAP00<br>8058 | 149.776542 |

Genome Annotated OBP
| Serial<br>No. | *An.culicifacies*<br>Transcript ID | Name | Signal<br>Peptide | OBP<br>Subfamily | Ortholog in<br>*An.gambiae* |
| --- | --- | --- | --- | --- | --- |
| 1. | ACUA027288 | OBP11 | Yes | Classic | AGAP002025 |
| 2. | ACUA007455 * | OBP64 | Yes | Classic | AGAP012324 |
| 3. | ACUA019075 | OBP18 | Yes | Classic | AGAP012319 |
| 4. | ACUA007086 * | OBP62 | Yes | Classic | AGAP002556 |
| 5. | ACUA016883 | OBP13 | Yes | Classic | AGAP002905 |
| 6. | ACUA005593 | OBP23 | Yes | Classic | AGAP012318 |
| 7. | ACUA011194 | OBP21 | Yes | Classic | AGAP008398 |
| 8. | ACUA019079 | OBP12 | Yes | Classic | AGAP002188 |
| 9. | ACUA022547 | OBP8 | No | Classic | AGAP000279 |
| 10. | ACUA009276 | OBP27 | Yes | Classic | AGAP012323 |
| 11. | ACUA018051 | OBP22 | No | Classic | AGAP010409 |
| 12. | ACUA023151 * | OBP60 | No | Plus-C | AGAP007281 |
| 13. | ACUA006032 | GOBP70 | Yes | Plus-C | AGAP006368 |
| 14. | ACUA005261 | GOBP68 | Yes | Plus-C | AGAP012658 |
| 15. | ACUA013374 | GOBP67 | Yes | Plus-C | AGAP007282 |
| 16. | ACUA003780 * | OBP44 | Yes | Two Domain | AGAP010648 |
| 17. | ACUA011054 * | OBP31 | Yes | Two Domain | AGAP010649 |
| 18. | ACUA011785 | OBP45 | Yes | Two Domain | AGAP010650 |
| 19. | ACUA011802 * | OBP35 | Yes | Two Domain | AGAP000642 |
| 20. | ACUA004895 | OBP42 | Yes | Two Domain | AGAP009065 |
| 21. | ACUA010086 * | OBP36 | Yes | Two Domain | AGAP000643 |
| 22. | ACUA020837 | OBP32 | Yes | Two Domain | AGAP000640 |
| 23. | ACUA028423 * | OBP40 | Yes | Two Domain | AGAP002191 |
| 24. | ACUA008136 * | OBP38 | No | Two Domain | AGAP000580 |
| 25. | ACUA017746 * | OBP39 | Yes | Two Domain | AGAP002190 |
| 26. | ACUA003948 | OBP41 | Yes | Two Domain | AGAP005182 |
| 27. | ACUA016063 * | OBP37 | No | Two Domain | AGAP000641 |

**\*Newly annotated OBPs.**

**Supplemental Table S4:** Catalogue of Olfactory Receptor.

| **Sl No .** | **Gene Accession No.** | **Receptor Type** | ***An. culicifacies* ID** | ***An. gambiae* Identity** | **CDS Origin** | **FPKM** |
| --- | --- | --- | --- | --- | --- | --- |
| 1. | XP_310061 | Or29 | ACUA025107 | AGAP009111 | Naïve_Transcript_6279 | 26.602643 |
| 2. | XP_315068 | OR35 | ACUA009020 | AGAP004971 | Naïve_Transcript_7874 | 11.9370834 |
| 3. | XP_315072 | OR31 | ACUA014252 | AGAP004974 | Naïve_Transcript_4545 | 27.0352063 |
| 4. | XP_309205 | OR36 | ACUA003785 | AGAP001012 | Naïve_Transcript_3706 | 14.6902145 |
| 5. | XP_311894 | OR62 | ACUA008089 | AGAP011978 | Naïve_Transcript_4436 | 17.7898333 |
| 6. | XP_313640 | OR26 | ACUA020087 | AGAP004357 | Naïve_Transcript_7012 | 19.519759 |
| 7. | XP_315773 | OR33 | ACUA020367 | AGAP005760 | Naïve_Transcript_6630 | 19.2215628 |
| 8. | XP_317124 | OR9 | ACUA008607 | AGAP008333 | Naïve_Transcript_6764 | 129.219612 |
| 9. | XP_321150 | Gustatory Receptor | ACUA025789 | AGAP001915 | Naïve_Transcript_6229 | 39.039518 |
| 10. | EFR27310 | IR21a | ACUA000154 | AGAP008511 | Naïve_Transcript_5794 | 8.28742772 |
| 11. | XP_311997 | IR41a | ACUA014728 | AGAP002904 | Naïve_Transcript_7850 | 23.907604 |
| 12. | XP_003436511 | IR41c | ACUA016051 | AGAP012951 | Naïve_Transcript_2012 | 1286.70859 |
| 13. | EFR22053 | Ionotropic Receptor NMDAR3 | ACUA007915 | AGAP005527 | Naïve_Transcript_5355 | 10.5566044 |
| 14. | XP_312117 | GLURIIc Ionotropic Receptor | ACUA017915 | AGAP002797 | Naïve_Transcript_6513 | 32.6012782 |
| 15. | XP_312026 | Putative GPCR class Orphan Receptor | ACUA015630 | AGAP002886 | Naive_Transcript_6435 | 17.9142377 |
| 16. | XP_311816 | OR45 | ACUA019417 | AGAP003053 | Naive_Transcript_2629 | 21.5803151 |
| 17. | XP_003436373 | Odorant Receptor | ACUA019417 | AGAP013396 | Naive_Transcript_7336 | 13.7599877 |
| 18. | XP_318786 | OR72 | ACUA020212 | AGAP009718 | Naive_Transcript_4446 | 7.45172072 |
| 19. | AGS08024 | OR41 | ACUA014638 | AGAP000226 | Naive_Transcript_5372 | 39.3789123 |
| 20. | XP_310066 | OR16 | ACUA017005 | AGAP009394 | 30min PBM_Transcript_4012 | 22.9711849 |
| 21. | XP_312289 | OR 39 | ACUA018263 | AGAP002639 | 30min PBM_Transcript_1117 | 151.09935 |
| 22. | XP_315048 | OR32 | ACUA026067 | AGAP004951 | 30min PBM_Transcript_8291 | 46.7776857 |
| 23. | XP_320543 | Or63 | ACUA028174 | AGAP011989 | 30min PBM_Transcript_7504 | 13.5053686 |
| 24. | XP_321007 | Or77 | ACUA025905 | AGAP002044 | 30min PBM_Transcript_3511 | 20.4188311 |
| 25. | XP_310173 | Or2 | ACUA019491 | AGAP009519 | 30min PBM_Transcript_6920 | 25.7922077 |
| 26. | XP_320128 | Putative GPCR class Orphan Receptor 16 | ACUA001535 | AGAP012427 | 30min PBM_Transcript_5058 | 9.13140271 |
| 27. | XP_308379 | IR75k | ACUA022596 | AGAP007498 | 30min PBM_Transcript_6656 | 9.88007955 |
| 28. | XP_003435724 | IR75I | ACUA008736 | AGAP005466 | 30min PBM_Transcript_5250 | 41.6081841 |
| 29. | EFR28910 | Or23 | ACUA005690 | AGAP007797 | 30min PBM_Transcript_7463 | 19.515343 |
| 30. | XP_314478 | Or44 | ACUA017149 | AGAP010505 | 30hr PBM_Transcript_8297 | 3270.8534 |
| 31. | XP_320541 | Or61 | ACUA01952<br>3 | AGAP01199<br>1 | 30hr<br>PBM_Transcript_3929 | 16.227537<br>7 |
| 32. | EFR26255 | Putative<br>GPCR<br>class an<br>orphan<br>receptor<br>21 | ACUA00174<br>9 | AGAP00568<br>1 | 30hr<br>PBM_Transcript_5866 | 5731.4820<br>1 |
| 33. | XP_309588 | Putative<br>GPCR<br>class an<br>orphan<br>receptor<br>4 | ACUA02127<br>9 | AGAP00403<br>4 | 30hr<br>PBM_Transcript_8157 | 1696.8165<br>9 |
| 34. | XP_320564 | IR76b | ACUA01504<br>7 | AGAP01196<br>8 | 30hr<br>PBM_Transcript_4542 | 48.344539<br>3 |
| 35. | EFR26426 | IR76a | ACUA00135<br>7 | AGAP00492<br>3 | 30hr<br>PBM_Transcript_7611 | 21.738059<br>8 |
| **Common Receptor Genes expressed in all the experimental conditions.** |  |  |  |  |  |  |
| 36. | ACQ55870 | Or 45 | ACUA01941<br>7 | AGAP00305<br>3 | 30hr<br>PBM_Transcript_4029 | 19.020802<br>4 |
| 37. | XP_556129 | Gustatory<br>Receptor | ACUA02126<br>0 | AGAP00549<br>5 | Naive_Transcript_2080 | 7531.7311<br>8 |
| 38. | XP_320553 | Or62 | ACUA00808<br>9 | AGAP01197<br>8 | Naive_Transcript_1886 | 2125.3509<br>4 |
| 39. | ABK97614 | Gr24 | ACUA01049<br>0 | AGAP00191<br>5 | Naive_Transcript_7268 | 661.57491<br>4 |
| 40. | XP_312379 | Orco | ACUA00346<br>9 | AGAP00256<br>0 | Naive_Transcript_972 | 425.79038<br>9 |
| 41. | XP_320874 | Or11 | ACUA01307<br>3 | AGAP01163<br>1 | Naive_Transcript_3225 | 341.52041<br>7 |
| 42. | XP_319142 | Gr22 | ACUA00405<br>4 | AGAP00999<br>9 | Naive_Transcript_1868 | 148.73984<br>9 |
| 43. | ACS83758 | Or66 | ACUA00031<br>6 | AGAP00331<br>0 | Naive_Transcript_5257 | 66.506607<br>4 |
| 44. | XP_312203 | Or28 | ACUA00979<br>6 | AGAP00272<br>2 | Naive_Transcript_2977 | 47.830094<br>4 |
| 45. | XP_314480 | Or24 | ACUA01191<br>5 | AGAP01050<br>7 | Naive_Transcript_6099 | 78.408311<br>3 |
| 46. | XP_00168872<br>6 | Or33 | ACUA02036<br>7 | AGAP00576<br>0 | Naive_Transcript_3900 | 25.280289<br>4 |
| 47. | XP_307763 | Gr17 | ACUA01976<br>7 | AGAP00325<br>5 | Naive_Transcript_7167 | 25.232859<br>2 |
| 48. | XP_319861 | Or29 | ACUA02510<br>7 | AGAP00911<br>1 | Naive_Transcript_5446 | 53.317533<br>4 |
| 49. | XP_321153 | Or8 | ACUA01438<br>3 | AGAP00191<br>2 | 30minPBM_Transcript_17<br>95 | 259.15027<br>7 |
| 50. | XP_313200 | Or42 | ACUA02484<br>8 | AGAP00427<br>8 | Naive_Transcript_7008 | 18.125490<br>5 |

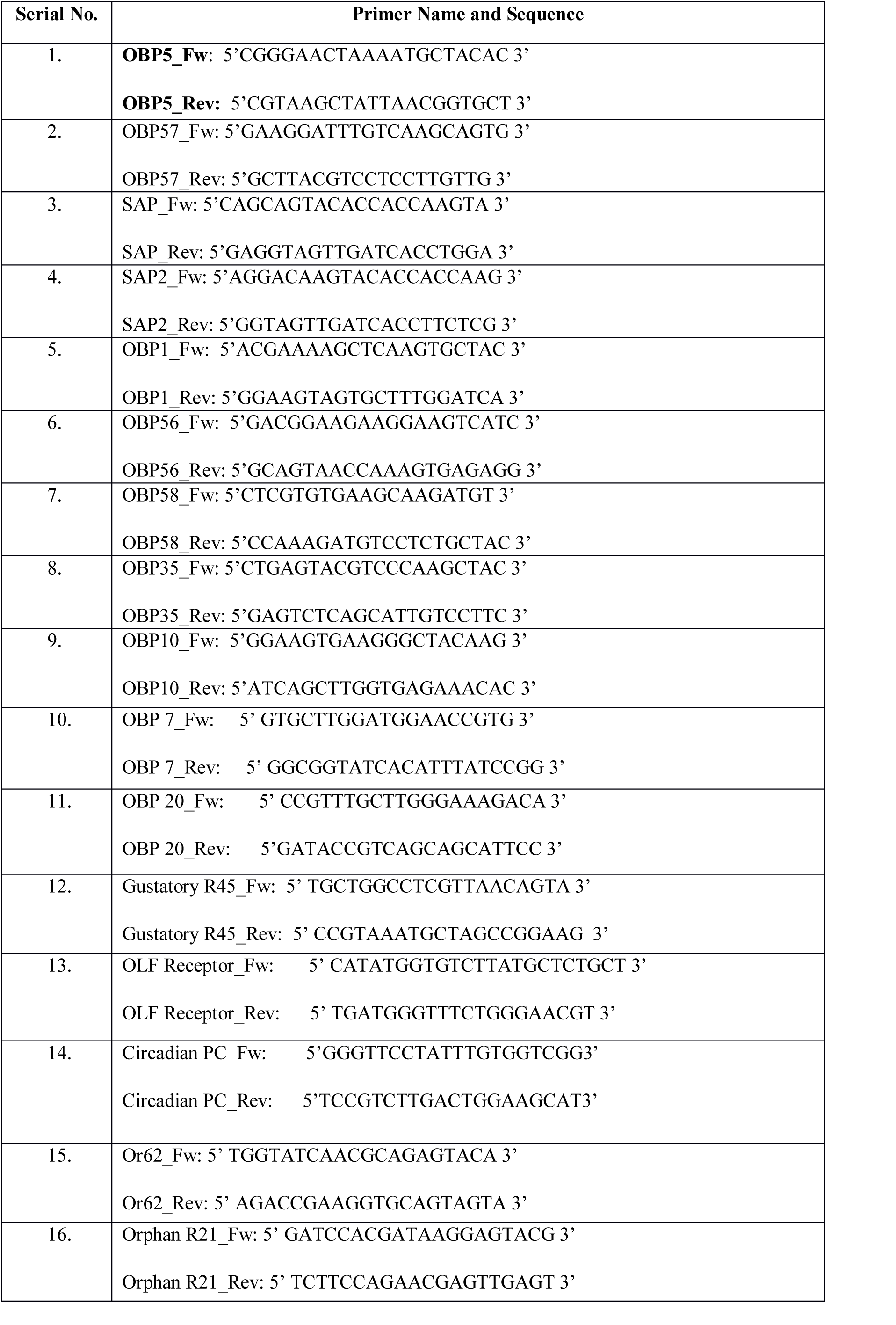
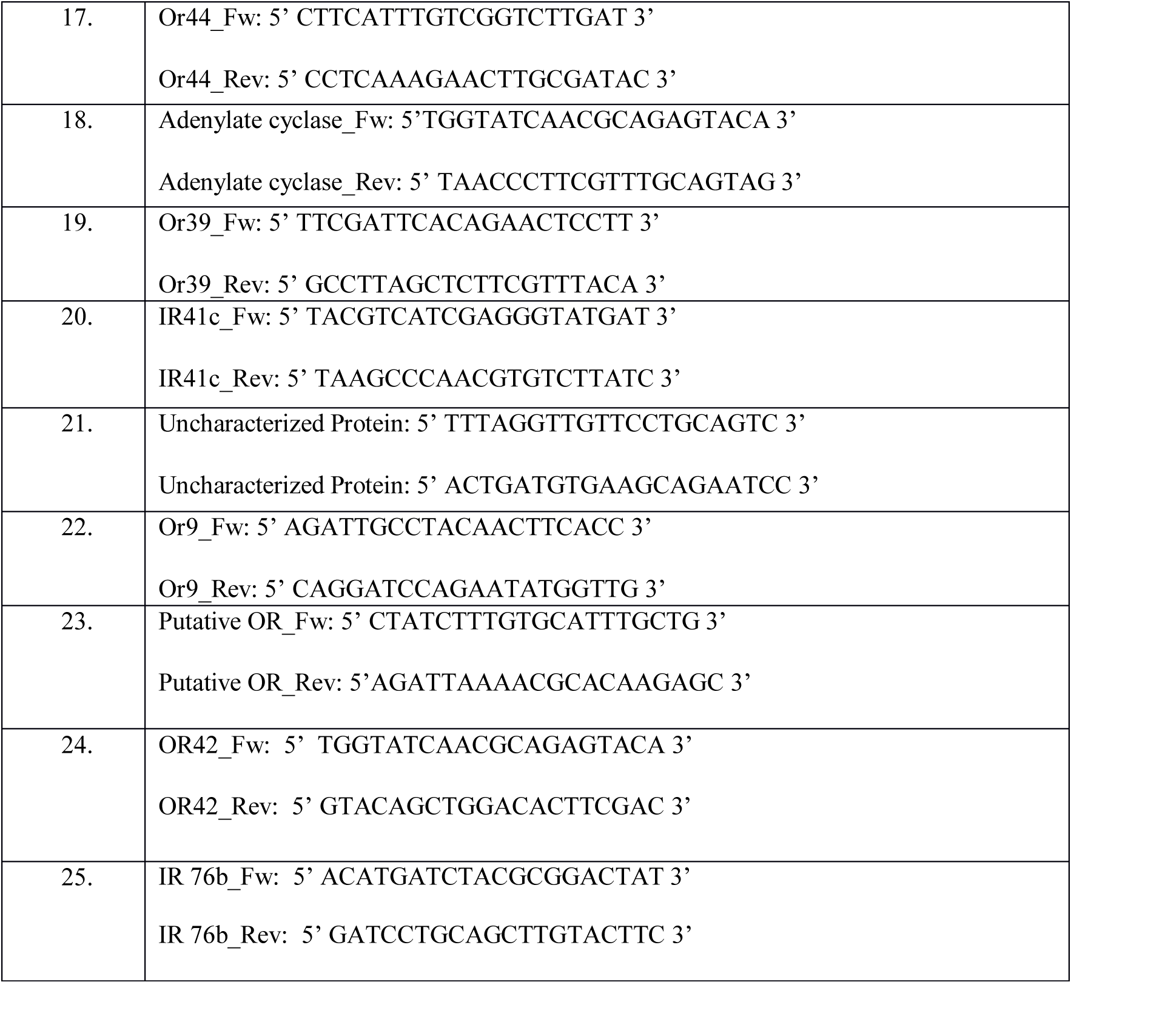
Primer Details.

**Table S6.** Showing major differences between *Anopheles culicifacies* and *Anopheles stephensi,* two major Indian Malaria vectors.

| ***Anopheles culicifacies*** | ***Anopheles stephensi*** |
| --- | --- |
| 1. ***Anopheles culicifacies*** contributes to about 60-65% of all malaria cases in India from rural to peri-urban areas and is widely distributed throughout the country.<br>2. **Species complexity:** *An. culicifacies* sp. comprised five sibling species provisionally designated as species A, B, C, D and E.<br>3. **Host Preference:** All the members of *An. culicifacies* are predominantly zoophilic except species E and rest indoor mainly in cattle sheds. Besides its low Anthropophagy, it acts as a major malaria vector due to the fact that it is found in high density.<br>4. **Breeding preference:** The preferred breeding sites are streams rice fields, seepage water borrow pits, irrigation channels, rain water collection etc.<br>5. **Biting Behaviour:** About 70–90% of *An. culicifacies* population caught during the whole night was found to feed prior to midnight during the months of January to April. Bimodal activity was seen during June and July indicating a further shift towards the second and third quarters of the night. During August–September, most biting takes place in the latter part of the night. | 1. ***Anopheles stephensi*** is an important vector of malaria in urban areas.<br>2. **Species complexity:** *An. stephensi* Liston 1901 lack sibling species complex, but exists as two forms, the type form and the variety *mysorensis*, which are distinguished by differences in the egg length and width and by the number of ridges on the egg float.<br>3. **Host Preferences:** In cities, it is the main malaria vector and has shown an increased tendency to feed on man than on cattle unlike in rural areas.<br>4. **Breeding preference:** *An. stephensi* is found predominantly in clear water, except in some polluted, blocked cemented drains with grass growth in the non-riverine zone near Delhi<br>5. **Biting Behaviour:** Biting occurs mostly before midnight and maximum activity was observed in the first quarter of the night (1800–2100 hrs). Biting activity has been observed till third and fourth quarters of the night, though at a low rate. *An. stephensi* was active throughout the night and 39% population fed before midnight. |

